# Expanding RNA editing toolkit using an IDR-based strategy

**DOI:** 10.1101/2023.12.23.573192

**Authors:** Minghui Di, Junjun Lv, Zhengyu Jing, Bryan Dickinson, Tian Chi

## Abstract

RNA base editors should ideally be free of immunogenicity, compact, efficient and specific, which has not been achieved for C>U editing. Here we first describe a compact C>U editor entirely of human origin, created by fusing the human C>U editing enzyme RESCUE-S to CIRTS, a tiny, human-originated programmable RNA binding domain. This editor, CIRTS-RESCUEv1 (V1), was inefficient. Remarkably, a short Histidine-Rich Domain (HRD), which is derived from the Internal Disordered Region (IDR) in the human CYCT1, a protein capable of liquid-liquid phase separation (LLPS), enhanced V1 editing at on-targets as well as off-targets, the latter effect being minor. The V1-HRD fusion protein formed puncta characteristic of LLPS, and various other IDRs (but not an LLPS-impaired mutant) could replace HRD to effectively induce puncta and potentiate V1, suggesting that the diverse domains acted via a common, LLPS-based mechanism. Importantly, the HRD fusion strategy was applicable to various other types of C>U RNA editors. Our study expands the RNA editing toolbox and showcases a general method for stimulating C>U RNA base editors.

## INTRODUCTION

Programmable RNA base editing is an important complement to DNA base editing (Rees and Liu, 2018). In contrast to DNA editing, RNA editing is reversible, in that once the editing is discontinued, the edited RNA is destined to be replaced by the nascent, unedited counterpart during RNA turn-over. This reversibility makes RNA editing intrinsically safer than DNA editing. For this reason, RNA editing would be particularly useful for mutating WT proteins to manipulate their physiological function in patients, where DNA editing is typically inapplicable for safety or ethical reasons. As a case in point, David Liu and colleagues used a DNA base editor to mutate *Ctnn1b (*encoding *β*-catenin) in the mouse inner ear, which results in Wnt activation in the cochlear supporting cells, thus paving the way for treating deafness via sensory cell regeneration (Yeh et al., 2018). Unfortunately, this strategy is clinically inapplicable, because the irreversible DNA editing would lead to persistent Wnt activation well known to be tumorigenic (Zhan et al., 2017). RNA editing, on the other hand, would fit the bill in such cases.

Two types of RNA editors have been described that convert A>I or C>U, as exemplified by the REPAIR and RESCUE platforms, respectively. The classical REPAIR platform comprises dCas13b fused to the deaminase domain of human ADAR2 (ADAR2dd), while the classical RESCUE platform comprises dCas13b fused to “RESCUE-S”, a bifunctional (A>I and C>U) deaminase domain derived from hADAR2dd via protein evolution and rational engineering (Abudayyeh et al., 2019; Cox et al., 2017; Liu et al., 2020). These classical editors have two major limitations.

First, the editors are too large to fit into AAV, thus hampering clinical applications. An attractive solution is to use engineered guide RNAs capable of recruiting endogenous ADARs to user-specified sites, thus bypassing the need for the REPAIR protein (Fukuda et al., 2017; Katrekar et al., 2022b; Merkle et al., 2019; Qu et al., 2019). However, this strategy is not applicable to C>U editing, because the endogenous C>U editing enzymes, in contrast to ADARs, have complex, stringent requirements for substrate conformation (Huang et al., 2020). Recently, Feng Zhang, Hui Yang and their respective colleagues have identified small Cas proteins (including EcCas6e and Cas13bt1) to facilitate AAV packaging, and successfully used them to upgrade the REPAIR and RESCUE platforms (Kannan et al., 2021; Wang et al., 2023; Xu et al., 2021). Unfortunately, these small Cas proteins cannot bypass the second limitation of the classic editors, which is common to all Cas-editors: the prokaryotic Cas proteins are potentially immunogenic in humans. Indeed, both innate and adaptive cellular immune responses to Cas9 have been detected in mouse models, and Cas9-specific antibodies and T lymphocytes are present in human plasma, which could compromise Cas9 therapeutic efficacy and also pose significant safety issues (Mehta and Merkel, 2020). The same caveat exists for other RNA editors utilizing prokaryotic RNA binding domains such as Lambda N and MCP (Katrekar et al., 2019; Montiel-Gonzalez et al., 2019; Montiel-González et al., 2016). The immunogenicity would be particularly worrisome for RNA editors, because the therapy might require repetitive editor delivery and antigen exposures, thus provoking prolonged/stronger immune responses.

Biological reactions can be facilitated by compartmentalizing the reactants into the membrane-less puncta/liquid droplets/condensates, which are formed via liquid-liquid phase separation (LLPS) driven by multi-valent weak interactions among Internal Disordered Regions (IDRs) in proteins (Alberti et al., 2019). IDRs are not only essential for LLPS, but also sufficient, because fusing IDRs to heterologous proteins such as GFP can confer the LLPS potential to the fusion partners. The IDRs, diversified in the primary sequence, have been characterized for numerous proteins. It is unclear whether IDRs can be harnessesd to boost RNA editors.

We have previously described CIRTS (Crispr/Cas Inspired RNA Targeting System), a tiny programmable RNA-targeting domain that is entirely of human origin (Rauch et al., 2019). We reasoned that CIRTS might be used to circumvent the aforementioned limitations of conventional RNA base editors. Here we report that CIRTS-RESCUE-S (version 1) was only a weak C>U editor but could be potentiated via IDR fusion (version 2), and this optimization strategy is applicable to various published C>U RNA base editors.

## RESULTS

### Development of CIRTS-RESCUE-S version 1 (V1)

V1 comprised RESCUE-S fused to CIRTS, the latter consisting of the human TAR Binding Protein (TBP) fused to human *β*-defensin (Fig. 1A, Editor #1); the TBP moiety recognizes the TAR hairpin in the gRNA, whereas *β*-defensin presumably acts by stabilizing the association of the editor with target transcripts and also by protecting the gRNA against nuclease attacks (Rauch et al., 2019). CIRTS can be recruited to target sites by a gRNA bearing the TAR hairpin linked to a targeting oligo (“spacer”) with a mismatched base opposite the targeted base (Fig. 1B) (Rauch et al., 2020, 2019). To facilitate the analysis of editing, we created a reporter construct expressing mouse *Ctnnb1* mRNA (Fig. 1C).

**Fig. 1.**
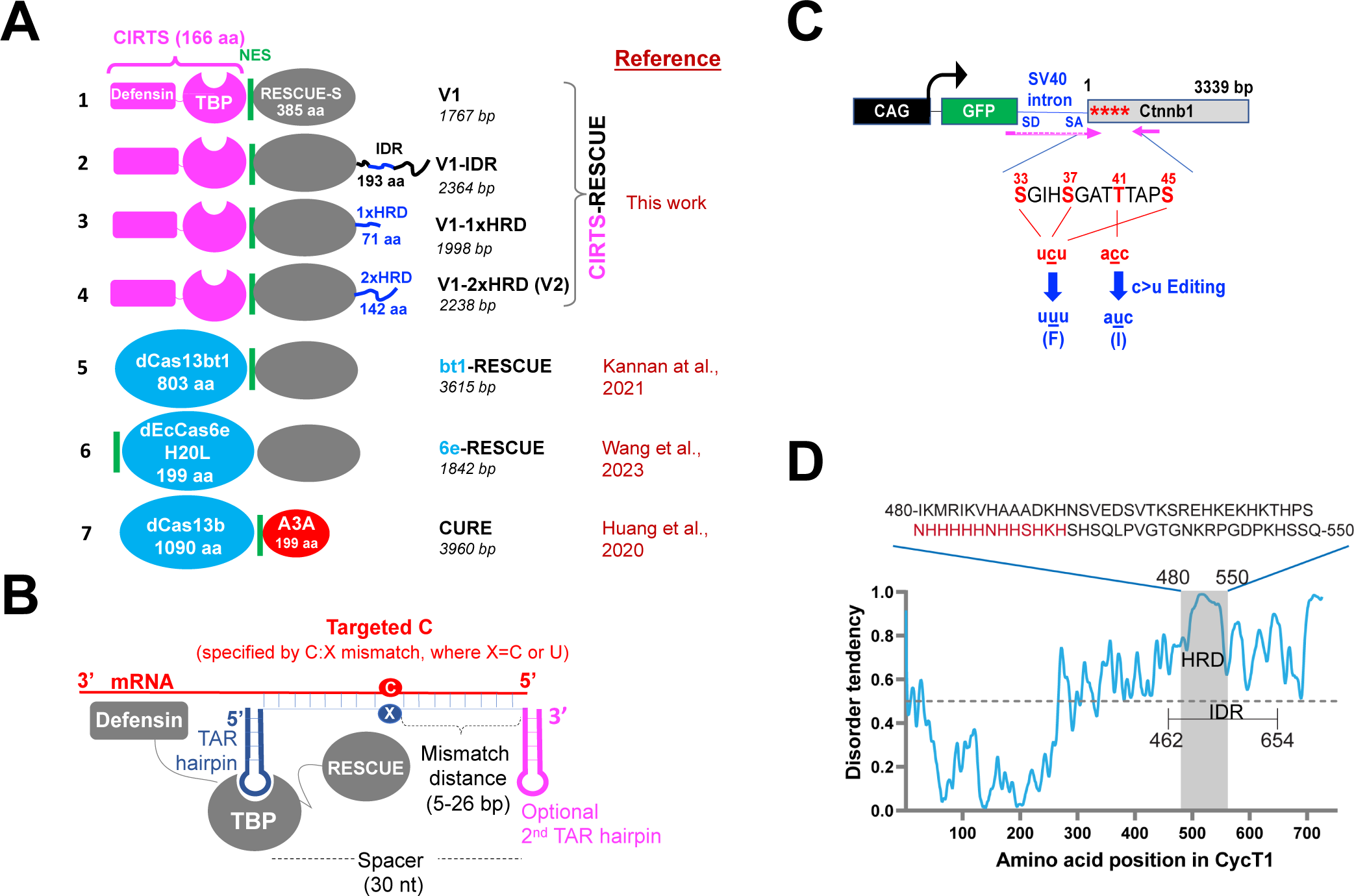
Experimental design. (A) The key editors tested in this study. RESCUE-S, the evolved bifunctional A and C deaminase domain. TBP, TAR-Binding Protein. CIRTS, CRISPR/Cas Inspired RNA Targeting System, a programmable RNA-binding domain comprising ý-defensin and TBP. IDR, Internal Disordered Region from CYCT1. HRD, Histidine Rich Domain within IDR. CURE, short-hand for CURE-C2, a version of the C-to-U RNA base editor comprising Apobec3A (A3A) deaminase fused to dCas13b. 6e-RESCUE is known as ceRBE in the original paper. (B) CIRTS-RESCUE gRNA configuration. The gRNA consists of a “spacer” fused to one or two copies of the hairpin of TAR, the HIV Trans-Activation Response element (thick blue or pink line). The spacer (thin blue line) is bound to the mRNA target sequence (red), whereas the hairpin associates with the TBP moiety of CIRTS-RESCUE. The target C on mRNA is specified by a mismatched base (X) at the spacer. The typical spacer length (30 nt) and mismatch distances (5-26 nt) are indicated; such gRNA configuration is designated 30/5-26. (C) *Ctnnb1* reporter. *Ctnnb1*, mouse *Ctnnb1* cDNA. Asterisks, phosphorylation sites required for CTNNB1 degradation, each of which can be mutated via C>U conversion at the mRNA level. SD and SA, splice donor and acceptor, respectively. Arrows, RT-PCR primers for amplifying mRNA, the forward primer spanning an SV40 intron inserted here to prevent the amplification of the *Ctnnb1* sequence carried in the contaminating plasmid. (D) Intrinsic disorder tendency of CYCT1. The central histidine cluster within the HRD is highlighted in red. Adapted from (Lu et al. 2018).

We first sought to optimize the gRNA. To this end, we tested the gRNAs carrying one or two copies of the TAR hairpin linked to a 30-nt spacer varied in the mismatched base (C or U) and the mismatch distance (5 to 26 nt, termed 30/5-26 hereafter; Fig. 1B). The gRNAs were co-expressed with V1-P2A-mCherry in the mouse neuroblastoma N2a cells for 48 hrs before the cells were analyzed for editing at *Ctnnb1* (S33F) at the reporter and endogenous mRNA (Fig. 2A). We found that at the reporter, dual TAR hairpins generally outperformed a single copy, and that the mismatch distance also impacted editing, with 15-26 nt much more effective than 5-10 nt, whereas the mismatch identity (C-or U-flip) was inconsequential (Fig. 2B, left). The same trend was seen at the endogenous transcript (Fig. 2B, right). Importantly, V1 editing efficiency was at most 35% and 24% at the reporter and endogenous mRNA, respectively, indicating that V1 was a weak editor.

**Fig. 2.**
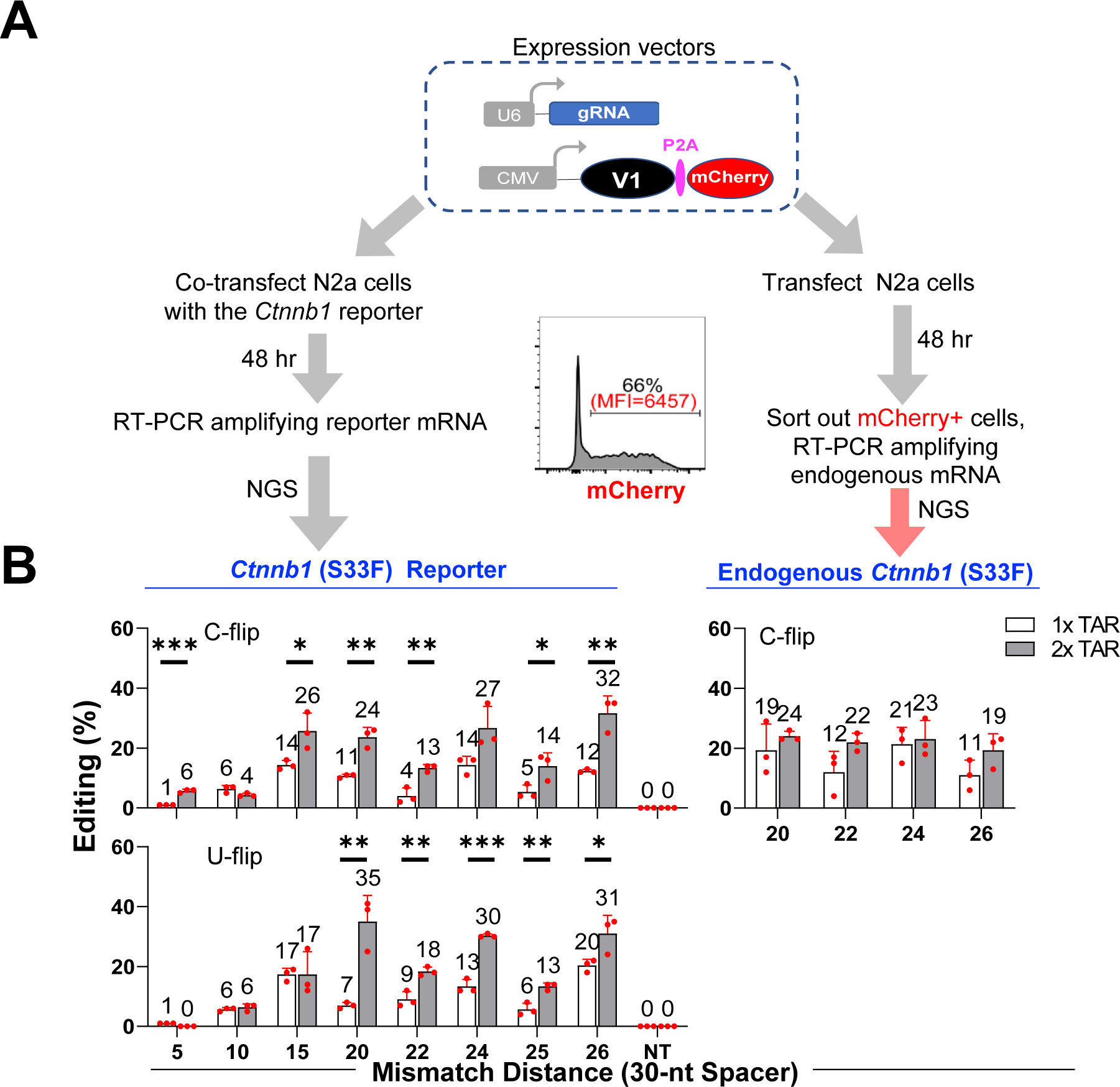
Boosting V1 editing via gRNA optimization. (A) Experimental workflow. P2A enables V1 and mCherry to be expressed from a single transcript. Shown also is a representative FACS plot of transfected N2a cells. Throughout this study, total cell population and the sorted mCherry+ subset were used for analyzing the editing (by NGS) at the reporter and endogenous transcripts, respectively. (B) Results from experiments testing the effects of hairpin copy number (1xTAR vs. 2x TAR) and gRNA mismatch distance on S33F conversion at the reporter (left) and endogenous (right) *Ctnnb1* mRNA. The nature of mismatch (C-flip vs U-flip) was also compared at the reporter. NT, gRNA bearing a random, non-targeting spacer. Unless otherwise noted, in all bar graphs in this study, values are mean+/-SD (n=3) and *P* values calculated using two-tailed Student’s t-test (\**P* ≤0.05, \*\**P* ≤ 0.01, \*\*\**P*≤ 0.001).

### Development of CIRTS-RESCUE-S version 2 (V2)

We attempted to potentiate V1 by fusing it to a small peptide capable of undergoing LLPS. We focused on the CYCT1 IDR because its core, the histidine-rich domain (HRD), known to be necessary and potentially sufficient for LLPS, comprises only 71 residue, one of the smallest LLPS domains characterized (Fig. 1A, Editor #2; Fig. 1D) (Lu et al., 2018). We first sought to test the effect of the IDR by fusing it to V1. Unexpectedly, the plasmid expressing the V1-IDR fusion protein proved difficult to construct, apparently due to IDR toxicity to our strains of competent cells. To bypass the cloning problem, we used PCR to synthesize the CMV-V1-IDR fragment and co-transfected it into HEK293T cells together with a plasmid expressing GFP, the latter intended for marking successfully transfected cells. We then isolated GFP+ cells by FACS and analyzed editor expression by Western blotting, finding V1 severely depleted after fusion with the IDR (Fig. S1A). However, Western blotting was at best semi-quantitative in our hands, unable to accurately and reproducibly quantify expression, which was contrary to FACS analysis of fluorescent proteins. Thus, we tagged V1-IDR with mCherry, repeated the transfection, and measured the fusion protein abundance using FACS. We found that the IDR caused a 2.8x reduction (from 85% to 30%) in the frequency of the mCherry+ subset among the transfected (GFP+) population, concomitant with a 10x reduction in mCherry mean fluorescence intensity (MFI) in the remaining mCherry+ cells, which collectively amounted to a 28x decrease in V1 expression (Fig. 3A). Thus, the FACS analysis confirmed and extended the conclusion based on Western blotting. The poor expression of V1-IDR-mCherry was also verified using fluorescence microscopy (see further in Fig. S5A).

**Fig. 3.**
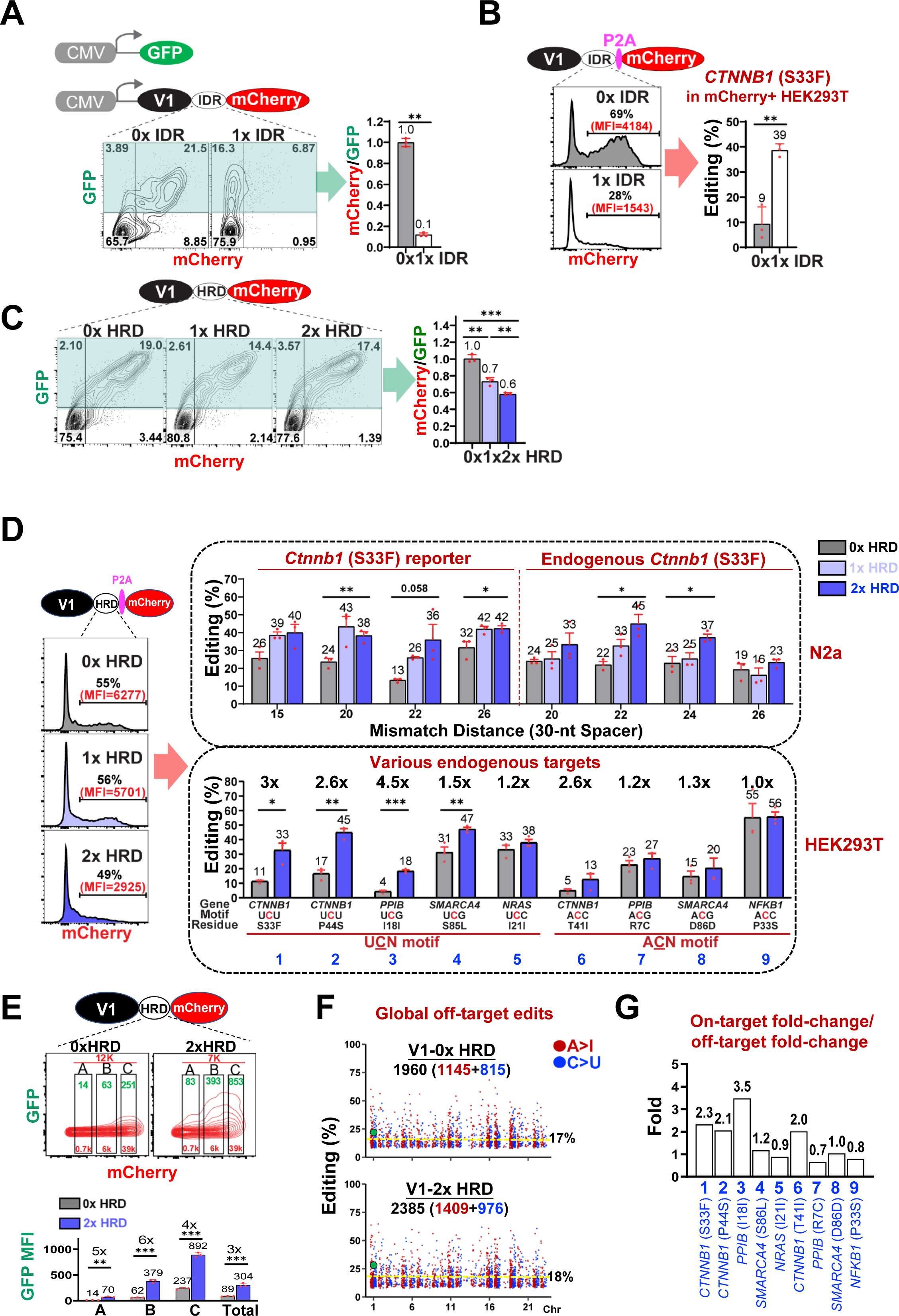
Development of V2. All the experiments were done in HEK293T cells unless otherwise noted. (A) The CYCT1 IDR dramatically reduced V1 expression. mCherry-tagged editors were co-expressed with GFP and examined by FACS 48 hrs after transfection. GFP marks successfully transfected cells (green shade) while mCherry reports editor abundance. The bar graph shows mCherry mean fluorescence intensity (MFI) in GFP+ population plotted relative to GFP MFI, with the value for V1 arbitrarily set as 1. (B) The CYCT1 IDR potentiated V1. DNA expressing editor-P2A-mCherry was transfected and mCherry+ cells sorted 48 hr later to quantify the S33F conversion at the endogenous *CTNNB1* mRNA. The gRNA configuration is 30/22 with C flip. (C) Same as A, but analyzing the effects of HRD (instead of IDR) on V1 expression. (D) Same as B , but analyzing the effects of HRD (instead of IDR) on V1 editing at *Ctnnb1* (S33F) in N2a cells (top bar graph) and a nine-target panel in HEK293 T cells (bottom bar graph). Representative FACS plots from an experiment in HEK293T cells are included. The gRNA configuration is 30/22 with U flip except that the gRNAs in the top bar graph have various mismatch distances as specified. (E) mCherry-tagged editors were co-transfected with a GFP reporter for C>U editing. The FACS plots are gated on total cells, where A, B and C mark arbitrary subpopulations with increasing mCherry expression (the red and green numbers are mCherry and GFP MFI, respectively). Data from triplicate transfections were summarized at the bottom. The gRNA configuration is 30/22 with C flip. (F) HRD mildly increased global off-target effects. Editor-P2A-mCherry was co-expressed in HEK293T cells with the gRNA targeting *NRAS* (I21I). The Manhattan plots display individual A>I (brown) and C>U (blue) off-target edits, together with edit numbers. Green dots, on-target edits; yellow lines, mean off-target editing rates. (G) Relative impact of 2x HRD on on-targets vs global off-targets. The fold-increases in the editing rates at various on-targets (Fig. 3D) are plotted relative to the mean fold-increase in the off-target editing (1.3x, Fig. 3F) .

We next determined the effect of the IDR on V1 editing. In theory, we could co-transfect the V1-IDR PCR fragment with the GFP expression vector and evaluate editing in GFP+ cells. However, the templates directing V1-IDR or GFP expression were not always co-transfected, thus confounding the analysis. To address this caveat, we inserted the “self-cleaving” P2A peptide between V1-IDR and mCherry, transfected the resultant V1-IDR-P2A-mCherry PCR fragment, and analyzed the mCherry+ cells; such cells by necessity co-expressed the V1-IDR cistron. Interestingly, the IDR moderately impaired mCherry expression even when P2A was inserted between the two proteins (Fig. 3B, left), perhaps because the V1-IDR cistron was poorly translated, causing a secondary defect in the translation of the mCherry cistron downstream. In any case, we sorted out mCherry+ cells and assessed editing at the endogenous *CTNNB1* mRNA. Remarkably, the IDR enhanced V1 editing 4x (from 9% to 39%; Fig. 3B, right). Given the much-reduced expression of V1-IDR relative to V1, the enhancement would likely be far more pronounced if estimated on a per-molecule basis.

Despite the powerful boosting effect of the CYCT1 IDR on V1 editing, its large size (193-aa) and the unintended repression of editor expression limited its application. We sought to bypass both problems using the core of IDR, namely the 71-aa HRD within the IDR (Fig. 1D)(Lu et al., 2018). In contrast to the IDR, the HRD proved non-toxic to the competent bacteria cells, enabling the construction of the plasmid expressing V1-HRD (Fig. 1A, Editor #3-4); for convenience, plasmids (instead of PCR fragments) were used to express various editors in all subsequent experiments except in Fig. 4A involving the CYCT1 IDR. Western blotting shows that HRD did not seem to compromise V1 expression (Fig. S1B), whereas FACS quantification of mCherry-tagged editor abundance revealed subtle and dose-dependent effects, with 1x and 2x HRD reducing relative V1 expression from 1 to 0.7 and 0.6 units, respectively (Fig. 3C).

**Fig. 4.**
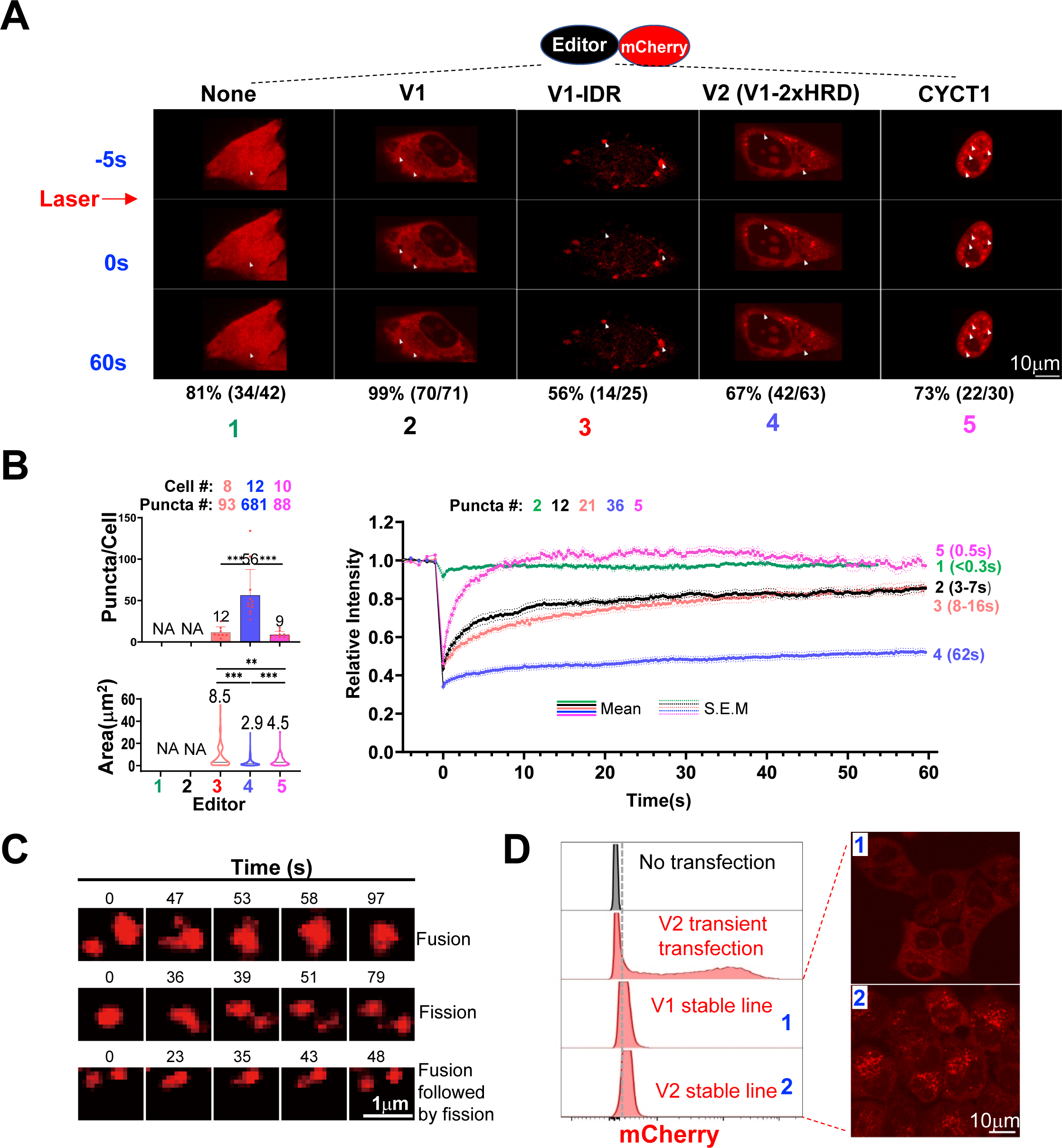
V1 potentiation is associated with LLPS. Experiments were done in HeLa cells. (A) The CYCT1 IDR/HRD induced LLPS in V1. Shown are representative images (100x) of the cells expressing indicated fluorescent proteins before (top) and after (middle and bottom) photo-beaching. The percentages of the cells with such images are displayed at the bottom, together with the actual numbers of the cells scored (bracketed fractions). Arrowheads indicate bleached spots. PCR products were used for transfection. (B) Same as A, but summarizing puncta abundance and size (left) in addition to FRAP data (right, bracketed numbers= **τ_1/2_**). NA, non-applicable. (C) Three representative forms of motion of the V2-mCherry puncta in a single cell imaged every second for 5 min. Images and the video of the whole cell are shown in Fig. S5C and Video S1, respectively. (D) V2-mCherry formed puncta even at minimal concentrations. HeLa cell lines with very low editor expression were analyzed by FACS and confocal microscopy in parallel. The cell image was digitally enhanced to visualize the faint puncta. FACS analysis of transiently transfected HeLa cells highly expressing V2-mCherry was included as a control (the second histogram).

Importantly, HRD markedly (up to 2x) stimulated V1 editing at *Ctnnb1* (S33F) in N2a cells both at the reporter and the endogenous mRNA, the effect rather robust to gRNA mismatch distances and partially dependent on HRD copy number (Fig. 3D, upper bar graph). V1-2x HRD was also able to edit other phosphorylation sites at the endogenous *Ctnnb1* (S37F, T41I, S45F; Fig. 1C), albeit less efficiently than S33F (Fig. S2). To determine whether the scenario at the *Ctnnb1* was generalizable to other targets and cell types, we compared V1 vs V1-2X HRD at nine endogenous target sites in the human HEK293T cells; these sites, bearing the U**<u>C</u>**N or A**<u>C</u>**N motifs, are known to be sensitive to RESCUE-S (whereas sites bearing the G/C**<u>C</u>**N motifs are refractory and thus not evaluated) (Abudayyeh et al., 2019; Huang et al., 2020). At all the nine sites except *NFKB1,* V1-2xHRD outperformed V1, where it increased the editing efficiency to various extents (up to 3x) at different sites, although the differences reached statistical significance in this particular trial at only 50% (4/8) of the sites (Fig. 3D, bottom bar graph). Of note, given that 2x HRD decreased V1 expression 1.7x (1/0.6; Fig. 3C), the potentiating effect of 2x HRD on V1 editing might be 1.7x higher on a per-molecule basis. To test this hypothesis, we took advantage of a reporter system we previously described, where C>U editing at the reporter mRNA leads to GFP induction, therefore enabling readout of editing efficiency as a function of editor abundance, if the editors are tagged with another fluorescent protein such as mCherry (Huang et al., 2020). Thus, we co-expressed the reporter mRNA and mCherry-tagged editors in HEK293T cells before FACS analysis 48hr post-transfection. Consistent with Fig. 3C, V1-mCherry expression exceeded V1-2x HRD-mCherry by 1.7x in the total transfected cell population (mCherry MFI being 12K vs 7K, Fig. 3E, FACS plots). The mCherry MFI, which varied substantially in individual transfected cells, indeed paralleled GFP MFI in the same cells (Fig. 3E, FACS plots), and when the former was normalized, V1-2x HRD-mCherry was found to induce 4-6x (∼5x) higher GFP MFI than V1-mCherry, contrary to only 3x when GFP in total mCherry+ cells was compared (Fig. 3E, bar graph, A-C vs Total). The data thus support our hypothesis that the activity of 2x HRD was underestimated due to reduced abundance of V1-2x HRD. Remarkably, the magnitude of underestimation (5/3 or 1.7x) coincided with the 1.7x reduction in editor abundance, attesting to the robustness of this reporter assay. Of note, direct mCherry fusion markedly compromised editor activity (Fig. S3). Thus, our standard approach throughout this paper is to use the direct mCherry fusion only for measuring editor expression, but split mCherry from the editors (via P2A) when assaying editing efficiency, and we infer per-molecule editing by normalizing the editing rate to the expression level.

Finally, we evaluated the effects of HRD on off-target editing globally at the transcriptome and locally at the bystanders, defined here as the Cs and As within the 60-nt region flanking the central targeted Cs. V1-2x HRD created 1.2x more global off-target edits than V1 (1690 vs 2385), concomitant with a subtle increase in the average editing rates (1.1x, from 17 to 18%), indicating that HRD exacerbated off-target editing by 1.2×1.1 or 1.3 fold (Fig. 3F). Importantly, this exacerbation (1.3x) was exceeded by the fold-increases (2.1-3.7x) in the on-target editing rates at five of the nine sites tested (Site #1, 2, 3, 4, 6), whereas the two were comparable at the remaining four sites, indicating that as a whole, HRD preferentially boosted editing at the on-targets over the global off-targets, thus increasing its relative specificity (Fig. 3G). In terms of bystander effects at the nine target sites, 2x HRD did not create any *de novo* edits, and had no or only limited effects on preexisting ones (Fig. S4). Specifically, 2x HRD had no effect on three sites lacking preexisting edits (#5, 7, 8) or one site with a single preexisting edit (#3). At the remaining five sites, V1 created a total of nine bystander edits, each of them clearly exacerbated by 2x HRD. Fortunately, V1 editing of these bystanders was generally very inefficient, at levels 6.7+/-4.9x below that of the respective on-targets, and so even after exacerbation by 2x HRD, the levels remained much lower than the respective on-target editing, differing by 4.6+/-3.8x (Fig. S4).

We conclude that 2xHRD boosted V1 editing at on-targets as well as off-targets, but the latter effect was comparably minor, indicating that on balance, 2x HRD had a beneficial effect. We have also tested 4xHRD, finding it inferior to 2x HRD in V1 potentiation (not shown). For simplicity, the upgraded version of V1 (V1-2x HRD) is termed V2 hereafter.

### Potentiated V1 form puncta *in vivo*

LLPS is characterized by microscopically visible puncta formation. To explore whether LLPS was implicated in V1 potentiation, we sought to image V1-CYCT1 IDR/HRD-mCherry in HEK293T and N2a cells. However, both cell types have scanty cytoplasm and, in our hand, tended to grow in aggregates instead of as a monolayer, thus hampering imaging. We therefore used HeLa cells free of such problems. We found that the positive control protein CYCT1-mCherry formed discrete nuclear puncta as previously reported (Lu et al., 2018), contrary to free mCherry or V1-mCherry proteins displaying diffuse fluorescence throughout the cell and in the cytoplasm, respectively (Fig. 4A, top row, Image 5 vs 1-2). These data confirmed that CYCT1 but not V1 underwent LLPS. In contrast to V1, V1-IDR-mCherry and V2-mCherry each formed discrete cytoplasmic puncta (Fig. 4A, top row, Image 3-4). Of note, in agreement with the Western blotting and FACS analyses, V1-IDR-mCherry fluorescence was extremely dim, necessitating digital enhancement for visualization (see Fig. S5A for the unadjusted image). On average, CYCT1, V1-IDR and V2 formed 9, 12 and 56 puncta per cell, respectively, with the V1-IDR puncta larger than the other two (8.5 vs. 2.9-4.5 *μ*m^2^ in mean size; Fig. 4B, left). Thus, V1 potentiation by the IDR and HRD was each associated with puncta induction.

We next characterized the mobilities of the mCherry-tagged molecules in the puncta using two assays. The first is Fluorescence Recovery After Photobleaching (FRAP). In this assay, a region of interest containing fluorescence-labeled molecules is bleached with a laser, and the fluorescence subsequently monitored as it recovers. We first sought to examine the free mCherry molecules not fused to other proteins. To this end, we illuminated a random fluorescent region for 0.3 s with a 561-nm wavelength laser, but found the fluorescence hardly diminished apparently due to extremely rapid exchanges of the bleached molecules with the surrounding unbleached ones (Fig. 4A, column 1; Fig. 4B, right, Curve 1). In contrast to free mCherry, this dose of illumination was sufficient to remove at least 50% of the fluorescence for various mCherry-tagged proteins, indicating that the fusion partners retarded mCherry mobility (Fig. 4A and Fig. 4B, right, Curve 2-5). Interestingly, the magnitude of retardation differed among the fusion proteins, as revealed by fluorescence recovery patterns: whereas CYCT1-mCherry fluorescence was restored fully and rapidly (within 10 s after bleaching), the restoration was slower for the three editors, reaching only ∼80% of the pre-bleach level for V1-(IDR) and ∼50% for V2 even at the end of the 60-second observation period (Fig. 4B, right). The recovery speed can be quantified using *τ*_1/2_, “the time for the exchange of half the mobile fraction between bleached and unbleached areas”, namely the time for achieving 50% of the total recovery (Ishikawa-Ankerhold et al., 2012). The *τ*_1/2_ was 0.5 s for CYCT1, 3-7 s for V1 and 8-16 s for V1-IDR (the fluorescence recovery for V1 and V1-IDR is assumed to max out at 80-100%, hence the ranges of *τ*_1/2_ values). *τ*_1/2_ for V2 was too long to determine within the 60-s observation window. We thus tracked the bleached V2 puncta for up to 10 min, finding that the recovery proceeded very slowly (*τ*_1/2_ =62 s) and maxed out at ∼70% of the original level (Fig. S5B). We conclude that the CYCT1 puncta were highly liquid-like, followed by the V1-IDR puncta, whereas the V2 puncta were the least mobile, but the biological significance of this observation awaits clarification (see Discussion). The second mobility assay is time-lapse microscopy, which revealed that V2 puncta underwent frequent fusion/fission that took place within minutes, consistent with the liquid-like behavior (Figs. 4C and S5C; Video S1). Similar fusion/fission events were observed for V1-1x HRD (Fig. S5D; Video S2).

IDR concentrations must be above threshold levels to initiate LLPS. As mentioned above, V1-IDR-mCherry was poorly expressed but still managed to undergo LLPS. Similarly, V2-mCherry formed discrete puncta even in the stable cell line expressing barely detectable levels of the editor (Fig. 4D). Such a strong propensity for LLPS may be particularly useful when editor expression is limiting, which might occur in some clinical applications.

Collectively, these data indicate that CYCT1 IDR and HRD induced puncta when fused to V1, thus providing the first line of evidence implicating LLPS in V1 potentiation.

### Distinct IDRs, but not an LLPS-impaired mutant, effectively replace the CYCT1 HRD to potentiate V1

To further explore the mechanisms of V1 potentiation, we evaluated four additional mammalian IDRs, derived from DDX4, DDX3X, FUS and HNRNPA1, respectively; these IDRs are all well-defined, and their sequences are unrelated (Fig. S6A) (Saito et al., 2019; Shin et al., 2017). We fused a single copy of these IDRs to V1, and compared their effects with that of 1x CYCT1 HRD. The IDRs from DDX4 and DDX3X behaved similarly to the CYCT1 HRD. Specifically, each IDR mildly decreased V1 expression (from 1 to 0.7∼0.8, Fig. 5A, Editor #1-4) but increased V1 editing at *Ctnnb1* S33F (from 50% to 61∼68%, Fig. 5B, Editor #1-4), which was associated with puncta formation (Fig. 5C, Editor #1-4). Interestingly, DDX3X harbors a 10-residue hydrophobic peptide (^37^RYIPPHLRNR^46^, Fig. S5A) potentially important for LLPS, as inferred from the role of a conserved peptide (^21^RYVPPHLRGG^30^) in LAF-1, the DDX3X paralog in the worm (Schuster et al., 2020). Indeed, deleting this peptide reduced V1-DDX3X IDR editing at *Ctnnb1* S33F (from 68% to 58%, *P*<0.001, Fig. 5B, Editor #4 vs 5), concomitant with a significant decrease in puncta number per cell (from 36 to 24, *P*<0.01, Fig. 5C, Editor #4 vs 5), which reinforces the role of LLPS in V1 potentiation.

**Fig. 5.**
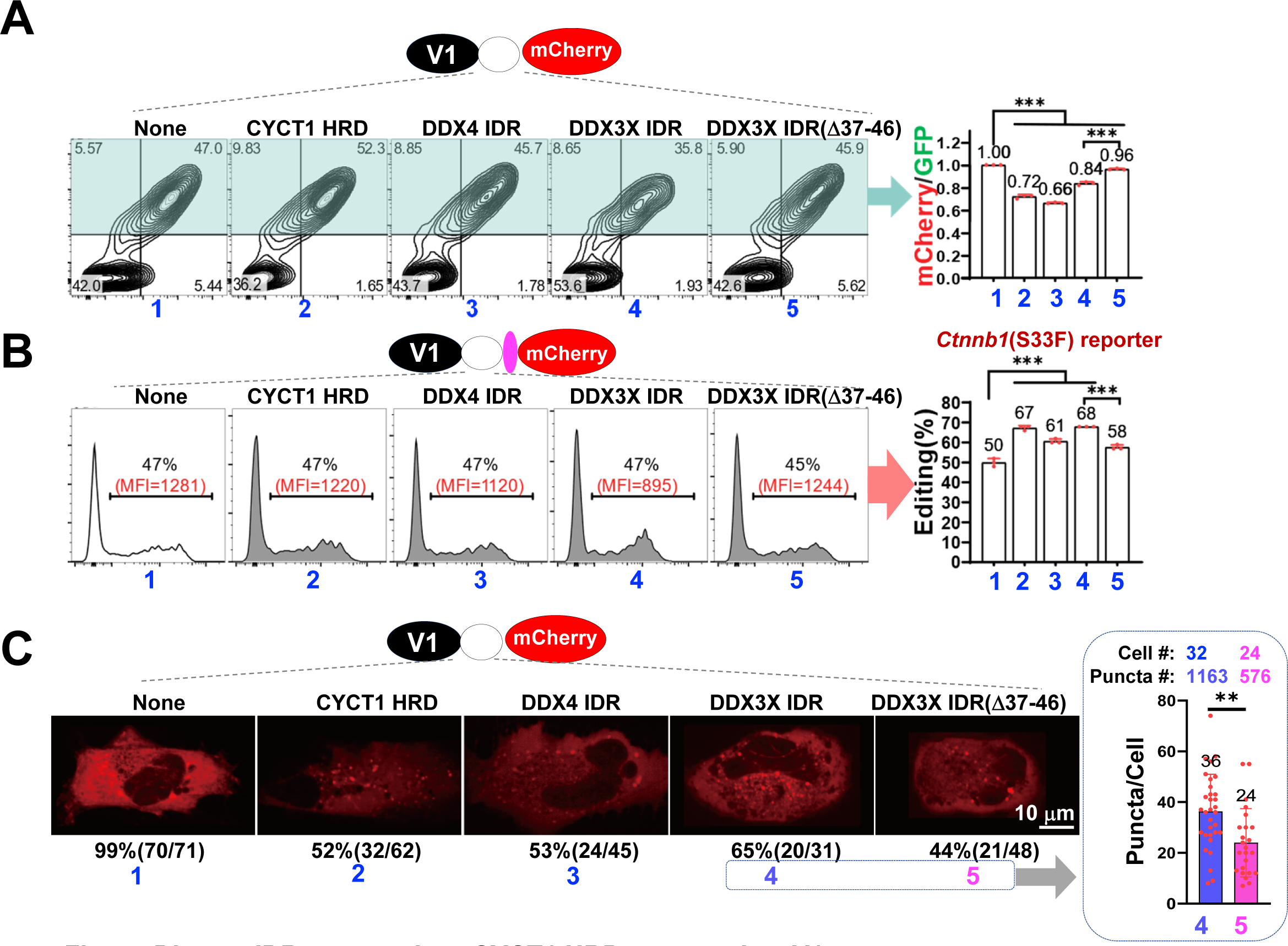
Diverse IDRs can replace CYCT1 HRD to potentiate V1. Shown are the expression (A) activity (B) and puncta formation (C) of the editors comprising V1 fused to indicated IDRs. Editor expression and editing were analyzed as in Fig. 3C and D, respectively, whereas puncta formation as in Fig. 4A. The bar graph in C summarizes the effect of DDX3X IDR deletion on puncta abundance. IDR sequences are listed in Fig. S6. The gRNA configuration in B is 30/22 with C flip.

In contrast to DDX4 and DDX3X, the FUS and HNRNPA1 IDRs proved inactive at enhancing V1 editing (Fig. S6B, left). The FUS IDR somehow failed to induce puncta, which can explain its inability to potentiate V1 (Fig. S6C, Image 3). On the other hand, V1-HNRNPA1 IDR did exist as puncta, but only in the nucleus (Fig. S6C, Image 4), apparently because this IDR harbors an NLS (Fig. S6A) that overwhelmed the NES in V1 (Fig. 1A) (Siomi and Dreyfuss, 1995). It is known that nuclear localization of RNA base editors can impair editing (Katrekar et al., 2019), which prompted us to compare V1-HNRNPA1 IDR with a nuclear version of V1. The nuclear V1 was indeed much weaker than V1 (6% vs 18% editing at the *Ctnnb1* reporter); remarkably, the HNRNPA1 IDR substantially enhanced editing by nuclear V1 (to 14%), just as 2x HRD (to 12%, Fig. S6B, right).

Thus, four unrelated LLPS domains (from CYCT1, DDX4, DDX3X and HNRNPA1) each potentiated V1, while a mutation in DDX3X IDR that impaired LLPS also compromised its potentiating effect, suggesting that the diverse LLPS domains acted via a common, LLPS-related mechanism.

### The CYCT1 HRD is applicable to various C>U editors beside V1

We next determined whether the IDR-based strategy is generally applicable to other types of C>U RNA base editors. We selected 2xHRD to address this issue, because among the LLPS domains tested, 2x HRD was relatively small and potent, and also able to induce LLPS even at extremely low concentrations. Specifically, we fused 2x HRD to three well-defined, representative RNA editors: dCas13bt1-RESCUE-S1 (“bt1-RESCUE”), dCas6e-RESCUE-S1 (“6e-RESCUE”) and CURE (Fig. 1A, Editor # 5-7). bt1-RESCUE is the prototypical small RNA editor derived from Cas13bt1, a compact version of the programmable RNA binding protein Cas13 (Kannan et al., 2021). 6e-RESCUE is also a small editor, but derived instead from Cas6e distinct from Cas13bt1 (Wang et al., 2023). Both editors use the bifunctional deaminase RESCUE-S to achieve C>U conversion, as in the case of V2. In contrast, CURE comprises the classic, large Cas13 protein (dCas13b) fused to Apobec 3A (A3A), a cytosine deaminase distinct from RESCUE-S (Huang et al., 2020). In contrast to RESCUE-S, A3A prefers the Cs located on the hairpin loop, and so CURE-mediated editing requires uniquely designed gRNAs capable of inducing a loop at the target site, contrary to the conventional gRNAs used by RESCUE-S-based editors. Thus, CURE is highly divergent from the other two editors in terms of structure and mechanisms of action. Remarkably, HRD stimulated all three editors (Fig. 6A), with little effect on their expression (Fig. S7). Specifically, bt1-RESCUE editing was boosted at all the nine targets (up to 2.3-fold at Site 3), as was 6e-RESCUE (up to 2.3-fold at Site 4), whereas CURE editing was enhanced at all the five sites bearing the U**<u>C</u>**N motif (Site 1-5, enhanced up to 4.5x at Site 1) albeit not at the remaining 4 sites bearing the A**<u>C</u>**N motif known to be refractory to CURE (Huang et al., 2020) (Fig. 6A).

**Fig. 6.**
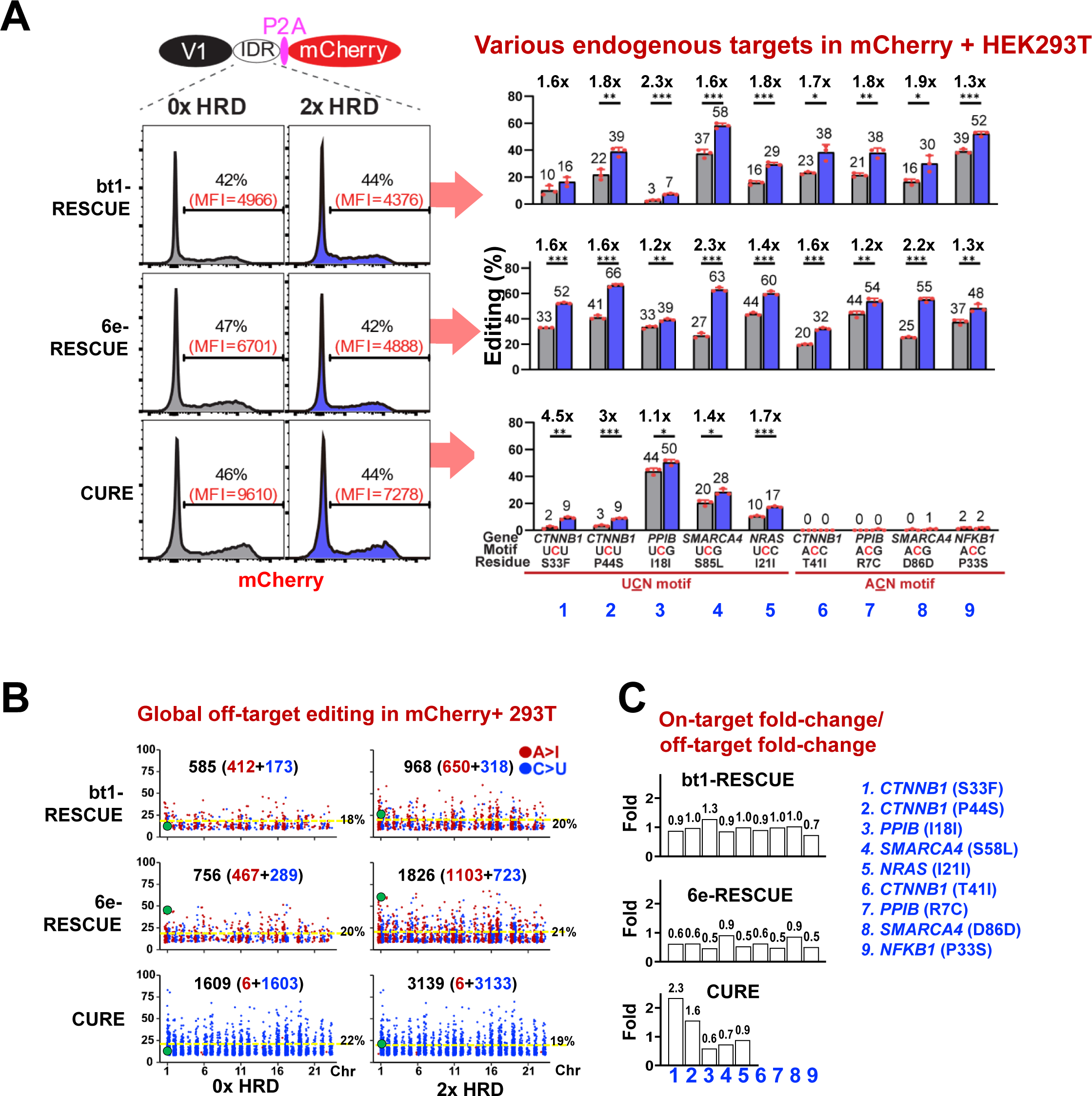
The CYCT1 HRD potentiates distinct RNA base editors in HEK293T cells. Shown are the effects of 2x HRD on the three representative published editors (# 5-7 in Fig. 1A) in terms of on-target editing (A) global off-target editing (B) and relative specificity (C) analogous to Fig. 3D, F and G, respectively; the panel of nine on-targets in A and C is the same as in Fig3. D and G. Optimal gRNA configurations are selected per original literature, which are 30/22 U flip, 30/26 C flip and S32L14 (32-nt spacer with 14-nt loop) for bt1-RESCUE, 6e-RESCUE and CURE, respectively.

2x HRD also exacerbated global off-target editing by bt1-RESCUE (1.9x), 6e-RESCUE (2.5x) and CURE (1.7x; Fig. 6B). In comparison with the effects of 2x HRD on on-target editing (Fig. 6A), the results revealed that, upon fusion of these editors to 2x HRD, the relative editing specificity was largely unaltered, compromised and enhanced/unaltered for bt1-RESCUE, 6e-RESCUE and CURE, respectively (Fig. 6C). Regarding bystanders, 2x HRD could not create *de novo* edits but tended to exacerbate preexisting editing, but the final editing levels were typically much below that of respective on-targets, just as in the case of V1 (Fig. S8). Remarkably, bt1-RESCUE and CURE were both highly specific, with bystander edits detectable at only one out of nine target sites (Site #6 and #2 for bt1-RESCUE and CURE, respectively), making both editors largely impervious to 2x HRD in terms of bystander effects (Fig. S8).

The data on the three representative editors, together with that on V1, show that 2xHRD increased on-target as well as off-target editing for all the four editors tested, with the off-target editing being predominant in only one (6e-RESCUE) of them, thus demonstrating the general utility of 2x HRD for RNA editor optimization.

## DISCUSSION

### An expended toolkit for RNA editing

This study introduced four new editors: V2 and enhanced versions of three preexisting editors (bt1-RESCUE, 6E-RESCUE and CURE), the latter created by fusing 2x HRD to the original versions of the editors. The HRD-induced increase in on-target editing was accompanied by the increase in off-target and bystander effects. Fortunately, in many cases we evaluated, the former increase was larger than, or at least comparable to, the latter. Thus, at these sites, the enhanced editors may be preferable over the original versions, especially when editing efficiency is limiting due to e.g., poor editor expression/delivery. The four new editors described in this study each have unique strengths and weaknesses, and are thus complementary in utility. For example, V2 is entirely human-originated and thus the only choice if host immunity is the major concern. V2 is also the smallest among all the four new editors, which adds to its utility. On the other hand, 6e-RESCUE-2x HRD is generally the most efficient, as first reported by Yang and colleagues (Wang et al., 2023) and confirmed by this study. It is also almost as small as V2, and thus generally the first choice when editing efficiency is the top priority. However, exceptions regarding editing efficiency were found at *NFKB1* and *PPIB I18I*, where V2 and CURE-2x HRD, respectively, were more active than 6e-RESCUE-2x HRD, and thus might be the top choice instead. Finally, bt1-RESCUE-2x HRD might prove the top performer at some other sites not tested in this work. Thus, occasionally, these editors may be preferable over 6e-RESCUE-2x HRD for maximal editing. We wish to emphasize that despite much effort, none of the editors available are perfect, and that development of an “ideal editor”, which must be at once free of immunogenicity, ultra-compact, highly efficient and specific, remains a key objective in the future. The IDR-based strategy, as detailed below, should help reach this objective.

### IDRs: utilities, mechanisms and limitations

The CYCT1 HRD potentiated four distinct C>U editors (V1, bt1-RESCUE, 6e-RESCUE and CURE); potentiated V1 formed puncta; distinct IDRs (DDX4, DDX3X, HNRNPA1) were able to replace the CYCT1 HRD to enhance editing and induce puncta; and finally, mutations in DDX3X IDR that impaired LLPS also compromised its ability to enhance editing. Collectively, these data indicate that IDRs can be harnessed to potentiate C>U editing, and suggest that the diverse domains worked via a common, LLPS-based mechanism, which is the working model embodying the parsimony principle, although further studies are necessary to conclusively exclude more complex hypotheses such as IDR-mediated recruitment of specific interaction partners.

Regarding the utility of IDRs in RNA editing, two issues are noteworthy. First, although we have focused on 2xHRD in this study, the full-length CYCT1 IDR was much more potent, as V1-1x CYCT1 IDR was as active as V1-2x HRD (namely V2; Fig. 3B vs 3D) even though 6x lower in expression (Fig. 3A vs 3C). However, the full-length CYCT1 IDR is not useful because it markedly enlarged and depleted the editors. Fortunately, numerous natural IDRs have been identified, and a library of designer minimalistic peptides (∼15-residues) has recently been reported that differ in LLPS propensity, dynamics and encapsulation efficiency (Baruch Leshem et al., 2023). These valuable resources can be systematically searched for the optimal LLPS domains/peptides. The parameters dictating the potency of the LLPS domains for stimulating RNA editing are unclear. The V1-1x CYCT1 IDR puncta were larger than V2 puncta (mean area 8.5 vs 2.9 *μ*m^2^), and also more dynamic: in the former puncta, fluorescence recovery after photobleaching proceeded faster (*τ*_1/2_=8-16 s vs 62 s) and plateaued at higher levels (>80% vs ∼70% of the initial intensity). Thus, large and highly mobile puncta seem correlated with higher editing efficiency, but this is a mere speculation at this point; indeed, the puncta formed by LLPS are known to differ widely in size, abundance and dynamics, but the physiological consequences of this variation remain elusive (McSwiggen et al., 2019). The second issue regarding the utility of IDRs is that the CYCT1 HRD (and by inference other IDRs as well) preferentially impacted on-targets over global off-targets, which is a pleasant finding. What might be the underlying mechanism? Presumably, concentrating an editor in the puncta can facilitate editing within the puncta, but at the same time jeopardize editing outside the puncta due to editor sequestration to the puncta. Thus, LLPS may have dual, conflicting effects on editing, which might differentially impact on-targets vs global off-targets. Specifically, the C>U editors used in this study are based on two deaminases, ADAR2 (V2, bt1-RESCUE and 6e-RESCUE) and A3A (CURE), whose substrates are cytosines embedded in dsRNAs and hairpin loops, respectively. For on-target editing, these secondary structures are induced *de novo*, after the gRNAs find and anneal to the target sequences, which may be a slow step that could be much accelerated inside the puncta; this acceleration may overweigh the negative impact of editor sequestration occurring outside the puncta, leading to a net increase in on-target editing. In contrast, for global off-target editing, the secondary structures of the substrates are preformed independently of gRNAs, leaving less room for acceleration by LLPS. As a result, the acceleration in off-target editing within the puncta might not markedly overweigh the negative effect outside the puncta, hence less conspicuous increases in off-target editing. Compared with global off-targets, bystander editing was more susceptible to exacerbation by 2x HRD, presumably because their editing required gRNA binding just as the on-targets. Importantly, 2x HRD could not create *de novo* bystander edits, and although it markedly exacerbated preexisting edits, such edits were rare and their basal editing rates typically very low, thus limiting the adverse impact of 2x HRD on the specificity of the editors.

Finally, we will discuss the limitations in the IDR-based approach and their countermeasures. First, as mentioned above, IDR can not only increase on-target editing, but also exacerbate off-target editing. Although in many cases, the former effect is predominant, it would still be highly desirable to minimize the latter, which might be achieved using a split-ADAR2 strategy capable of abolishing the global off-target edits (Katrekar et al., 2022a). Second, fusion of IDRs enlarges the editors, which may be acceptable for editors such as 6e-RESCUE (1.8 kb), but hamper AAV packaging for others editors such as bt1-RESCUE (3.6 kb, which already approaches the 4.7 kb size limit when linked to a gRNA expression cassette). Using the 15-residue minimalistic LLPS peptides or the split-ADAR2 strategy mentioned above can potentially solve the problem. Lastly, IDRs can potentiate CRISPRa (Liu et al., 2023; Ma et al., 2023) but unable to stimulate Cas13d, a programmable RNA-cutting enzyme (unpublished). Thus, we propose that the utility of IDR-based strategy in various editing platforms (including CRISPR editing and CRISPR interference) be carefully characterized on a case-by-case basis.

## METHODS

### Constructs

The plasmids were constructed using routine methods, while the templates expressing V1-CYCT1 IDR and its related/control proteins were synthesized by PCR due to cloning problems (Supplementary Information).

### Cell culture and transfection

The mouse neuroblastoma line N2a and human embryonic kidney line HEK293T (both from ATCC) were cultured at 37°C with 5% CO2 in DMEM containing high glucose, sodium pyruvate, penicillin–streptomycin and 10% fetal bovine serum. Cells were passaged three times per week and tested to exclude mycoplasma contamination. Transfections were performed with Lipofectamine 3000 in 48-well or 24-well plates per manufacturer’s instruction. Briefly, cells were plated into 48-well plates at 5 × 10^4^/well the first day and transfected 1 day later. DNA was mixed with 1*μ*L Lipofectamine P3000 (Thermo Fisher Scientific, L3000015) into 25 *μ*L Opti-MEM (Invitrogen) and incubated for 5 min at room temperature. 0.75 *μ*L of the Lipofectamine 3000 (Thermo Fisher Scientific, L3000015) was diluted into 25 *μ*L Opti-MEM (Invitrogen) and combined with the DNA: P3000 mixture, incubated for another 20 min at room temperature. The DNA: P3000: Lipofectamine 3000 mixture was added dropwise into the wells. Cells were analyzed ∼48 hr post-transfection.

### Editing at the *Ctnnb1* reporter and various endogenous transcripts

To analyze editing at the reporter, the *Ctnnb1* reporter plasmid (50 ng) was co-transfected into N2a or HEK293T cells in 48-well plates together with the vectors expressing editors (200 ng) and gRNAs (250 ng) or, where applicable, non-targeting (NT) gRNA bearing a random spacer sequence. 48 hr later, cells were harvested for RNA analysis as described (Joung et al., 2017). Briefly, 10 *μ*L lysis buffer was added per 5 × 10^4^ cells. After a 10-min incubation, 1 *μ*L stop solution was added before 1 *μ*L of the lysate was used for reverse transcription with HiScript II Q RT SuperMix kit (Vazyme, R223-01). The edited sites were PCR amplified before NGS analysis on the Illumina HiSeq platform or occasionally Sanger sequencing, as described previously (Huang et al., 2020). To assay editing at endogenous transcripts, N2a or HEK293T cells in 24-well plates were transfected with the vectors expressing editors (400 ng) and gRNAs (600 ng). 48 hr later, the mCherry+ cells were sorted and total RNA isolated using the NucleoZOL (Macherey-Nagel). Downstream steps were as described for the *Ctnnb1* reporter. The PCR primers are listed in Supplemental Information.

### Stable cell lines

To make stable lines for evaluating puncta formation at low protein concentrations (Fig. 6D), *piggy*Bac vector expressing V1/V2-mCherry (800 ng) was co-transfected with PBase expression vector (200 ng) into HeLa cells in 24-well plates. Single cells with barely detectable mCherry fluorescence were sorted 2 weeks later into 96-well plates, and clonally expanded for 2 weeks before imaging.

### Global off-targets at the transcriptome

The procedure was as described (Huang et al., 2020). Briefly, HEK293T cells in 6-well plates were transfected with the vectors expressing the *NRAS* gRNA (1.8 *μ*g) and editor-P2A-mCherry (1.2 *μ*g). mCherry+ cells were sorted 48 hr later and their total RNA isolated using NucleoZOL (Macherey-Nagel). The mRNA fraction was then enriched using a NEBNext Poly(A) mRNA Magnetic Isolation Module (NEB) before library construction using NEBNext Ultra RNA Library Prep Kit for Illumina (NEB). The libraries were sequenced on an Illumina NoveSeq 6000-PE150, at a depth of at least 6G per sample. The reads were mapped to the human reference genome (hg38) by STAR (2.7.1) using the 2-pass mode. Duplications were removed using Picard (2.20.5), and SNV identified by GATK HaplotypeCaller (4.1.3), with QD (Quality by Depth) > 2, sequencing coverage > 50× and reads in the wild-type samples supporting the reference allele > 99%. Edits shown in this study are intersection sets of two replicates. The editors in Fig. 3F and Fig. 6B were analyzed in parallel and the results thus directly comparable.

### Western blotting

To assess the effect of IDR on V1 expression, HEK293T cells in 6-well plates were co-transfected with 1.5 *μ*g PCR products expressing HA-tagged V1-IDR or V1 together with a plasmid expressing mCherry (0.5 *μ*g). mCherry+ cells were sorted 48 hr later, lysed and probed with HRP-conjugated antibodies against HA or β-actin (Cell Signaling Technology). Proteins were visualized with Omni-ECL™Enhanced Pico Light Chemiluminescence Kit (Epizyme) and imaged using the Amersham Imager 600 Chemiluminescent Imaging System (AI600). The effect of HRD on V1 expression was similarly analyzed except that editor-expressing plasmids instead of PCR products (2.7 *μ*g) were co-transfected with mCherry-expressing vector (0.3 *μ*g).

### Fluorescence imaging and FRAP

HeLa cells were transfected with DNA (1 ug) expressing mCherry-tagged proteins cultured on 29-mm glass-bottom dish (Cellvis); plasmids (1 ug) were used for transfection except in Fig. 4A, where PCR amplicons (1 ug) were used instead. At around 48 hours post-transfection, cells were observed utilizing a Nikon spinning disk microscope equipped with a dual-laser system. Imaging was conducted using a low-intensity 561-nanometer laser, while employing a 100x magnification objective with a numerical aperture of 1.35. For FRAP, the puncta were bleached by 300-ms illumination with a high-intensity 561-nm laser at 30% of the maximal intensity, which reduced fluorescence by ∼50%. Recovery of fluorescence into the bleached spot was imaged continuously with 300-ms exposure time for up to 10 min using a low-intensity laser; average pixel intensity in the bleached region was corrected for photobleaching caused by the low-intensity illumination, and plotted vs time.

### Statistical analysis

To analyze the editing efficiency and editor-mCherry expression, transfection was done typically in triplicates, and at least two biological repeats were performed. The relative differences among different editors compared within an experiment were highly consistent across different biological repeats, whereas the absolute values can vary (usually less than 2-fold) between the repeats. The bar graphs display the results from one of the biological repeats, where the values are mean+/-SD from triplicate transfections (as shown by the numbers of the dots). *P* values were calculated using two-tailed Student’s T-test (\**P* ≤0.05, \*\**P* ≤ 0.01, \*\*\**P*≤ 0.001). The experiments were not randomized and the investigators not blinded to allocation during experiments and outcome assessment.

## AUTHOR CONTRIBUTIONS

T.C. designed and supervised the project. B. D. provided guidance on CIRTS. Other authors performed the experiments .

## ACKNOWLEDGEMENT

We thank the Molecular and Cell Biology Core Facility (MCBCF) and the Molecular Imaging Core Facility (MICF) at the School of Life Science and Technology, ShanghaiTech University for providing technical support, Cong Liu for advice on LLPS, and Joyce Chi for proofreading. This study is supported in part by a grant to T.C. from National Science Foundation of China Major Research Plan (#92068115) and Shanghai Frontiers Science Center for Biomacromolecules and Precision Medicine at ShanghaiTech University.

---

**Fig. S1.**
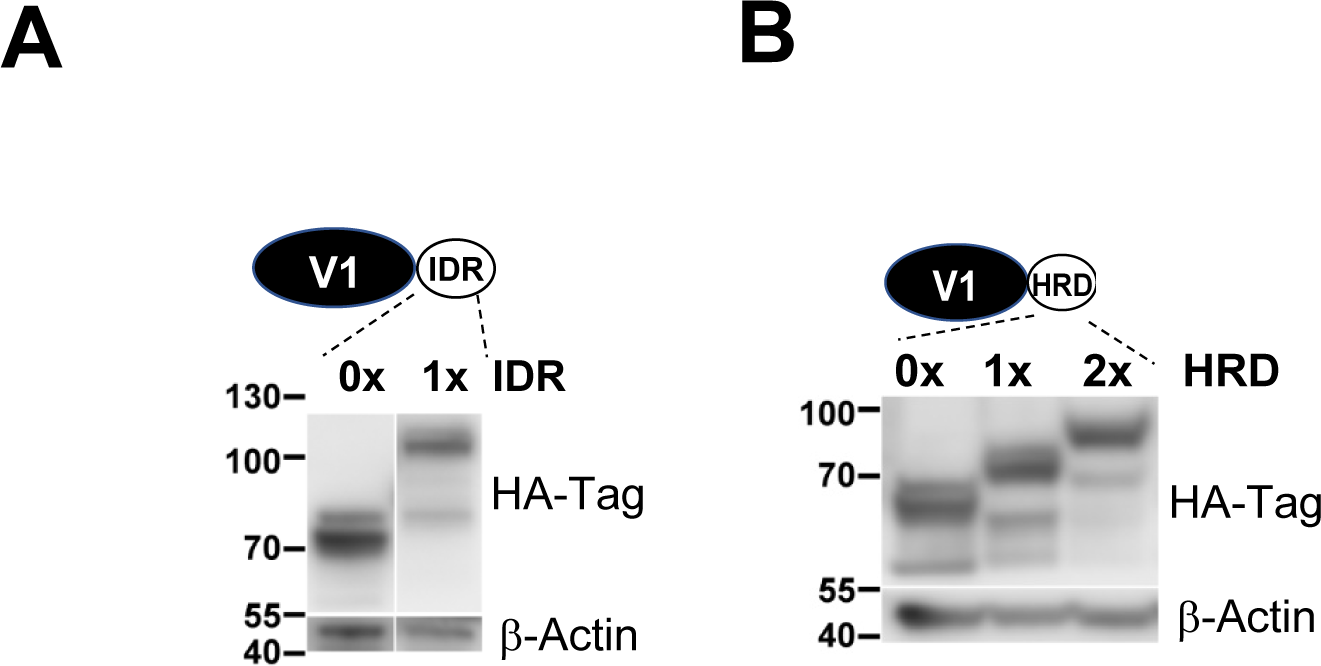
Western blotting analysis of V1-IDR (A) and V1-HRD (B) expression. HEK293T cells were transfected with 1 ug of PCR fragments (A) or plasmids (B) expressing HA-tagged editors, and probed with anti-HA and actin antibodies 48 hr post-transfection.

**Fig. S2.**
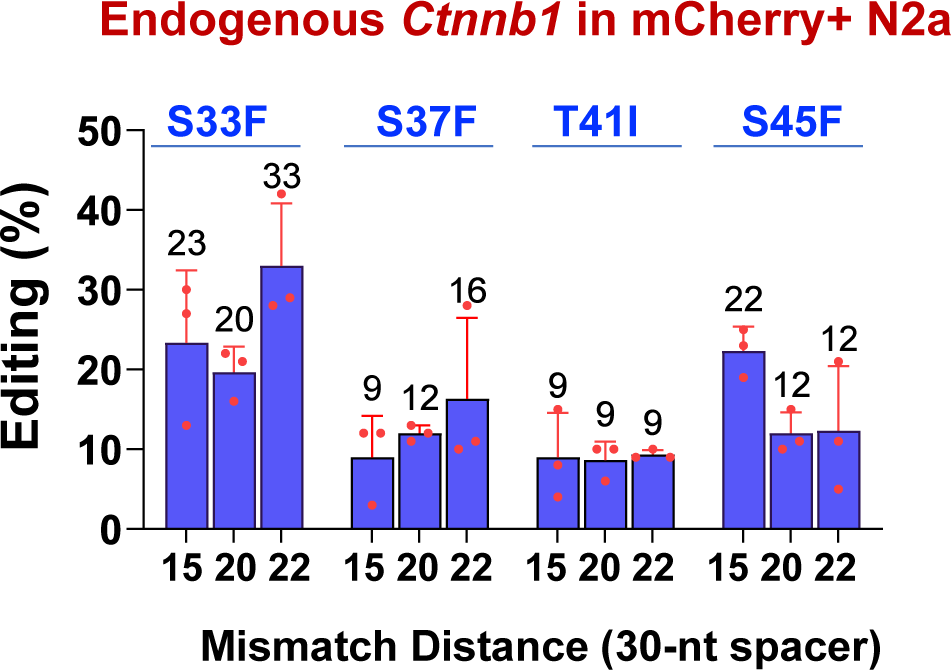
V2 is able to edit various phosphorylation sites at *Ctnnb1*. These sites are depicted in Fig. 1C. The experiment was done as in Fig. 3D, but only V2 and the indicated sites were tested.

**Fig. S3.**
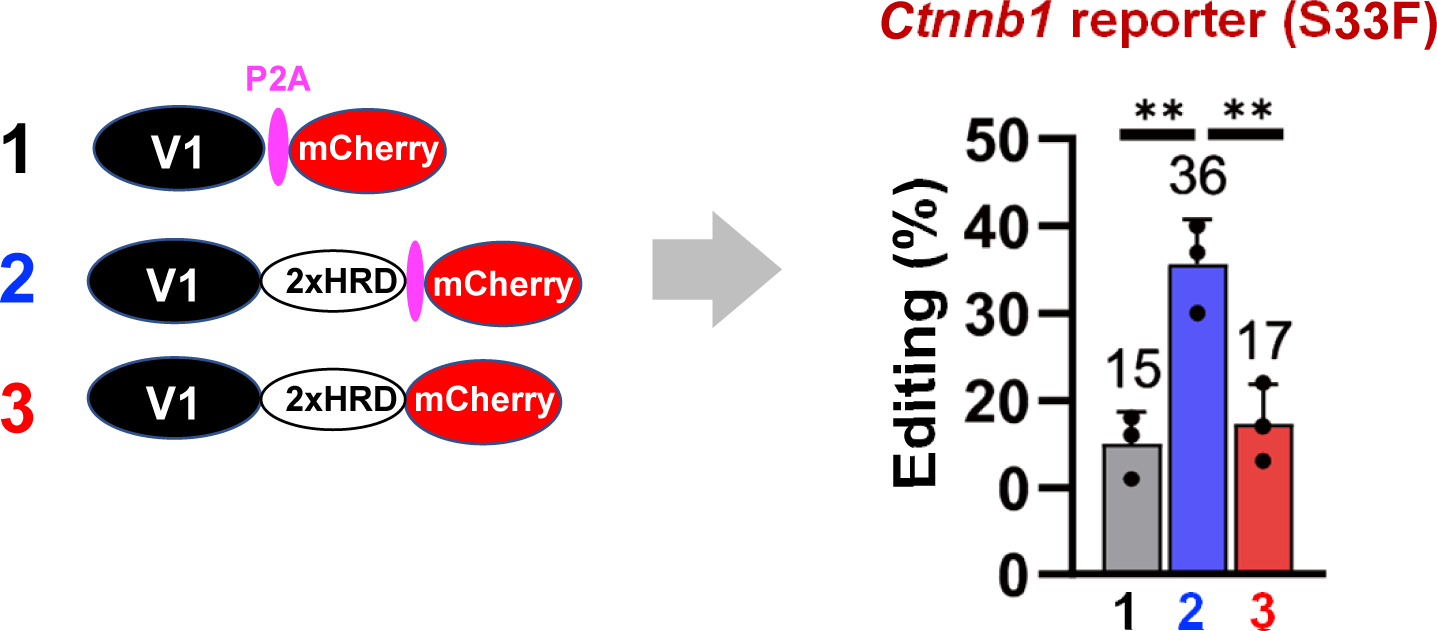
mCherry interferes with editing. 293T cells were transiently transfected with the plasmids expressing the editors diagrammed at the left, and editing analyzed 48hr later . The data show that direct fusion of mCherry leads to a 2x decrease in the editing efficiency (right, 2 vs 3).

**Fig. S4.**
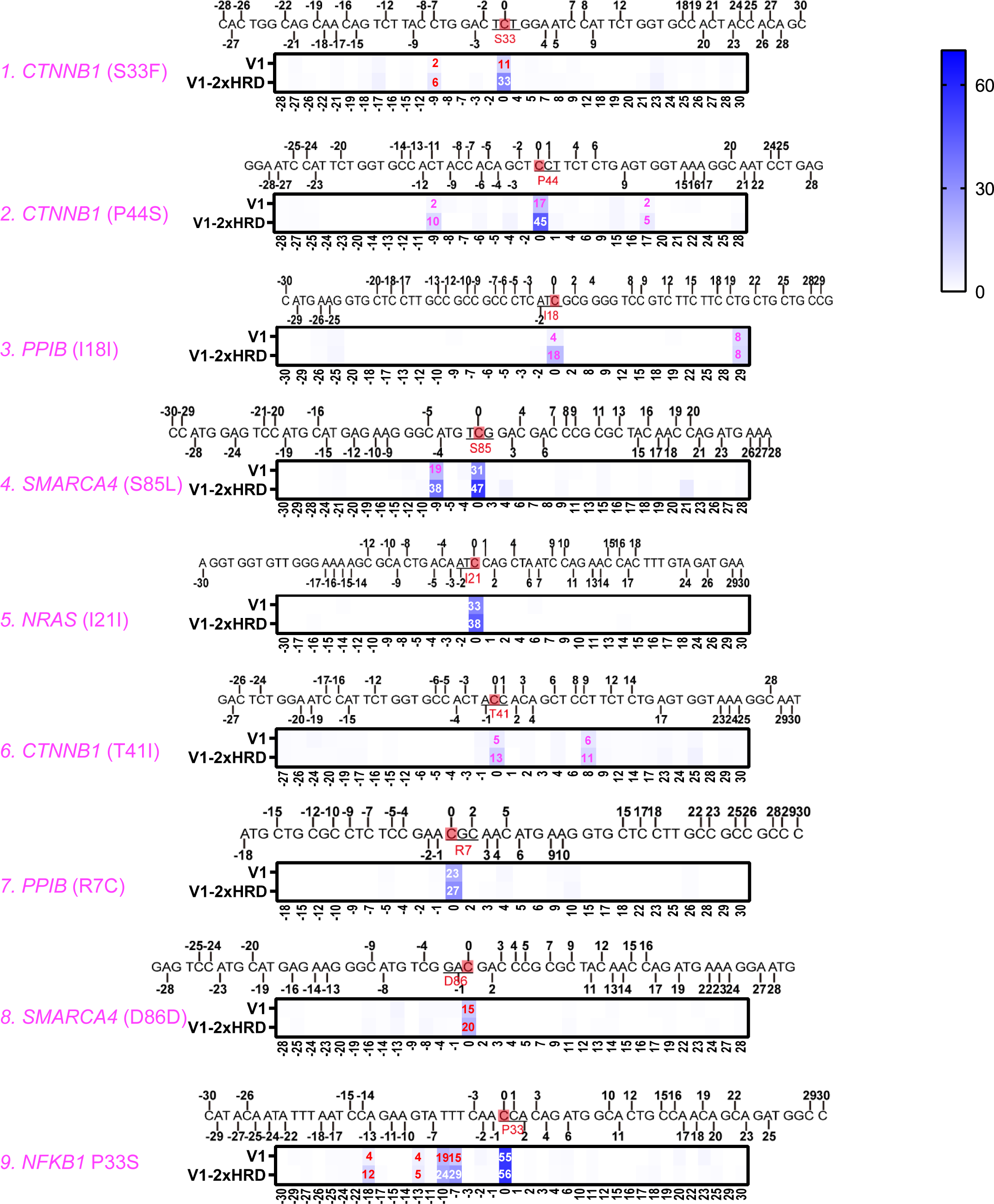
Effects of 2x HRD on bystander editing. Same experiment as in Fig. 3D (bottom), but showing the targeted Cs (red) together with the bystanders (numbered bases) within the 60-nt flanking regions. The values in the heatmap denote editing rates, and are displayed only for the bases clearly and reproducibly edited. Note that 2x HRD did not exacerbate the bystander effect at Site 3, as in Fig. S8.

**Fig. S5.**
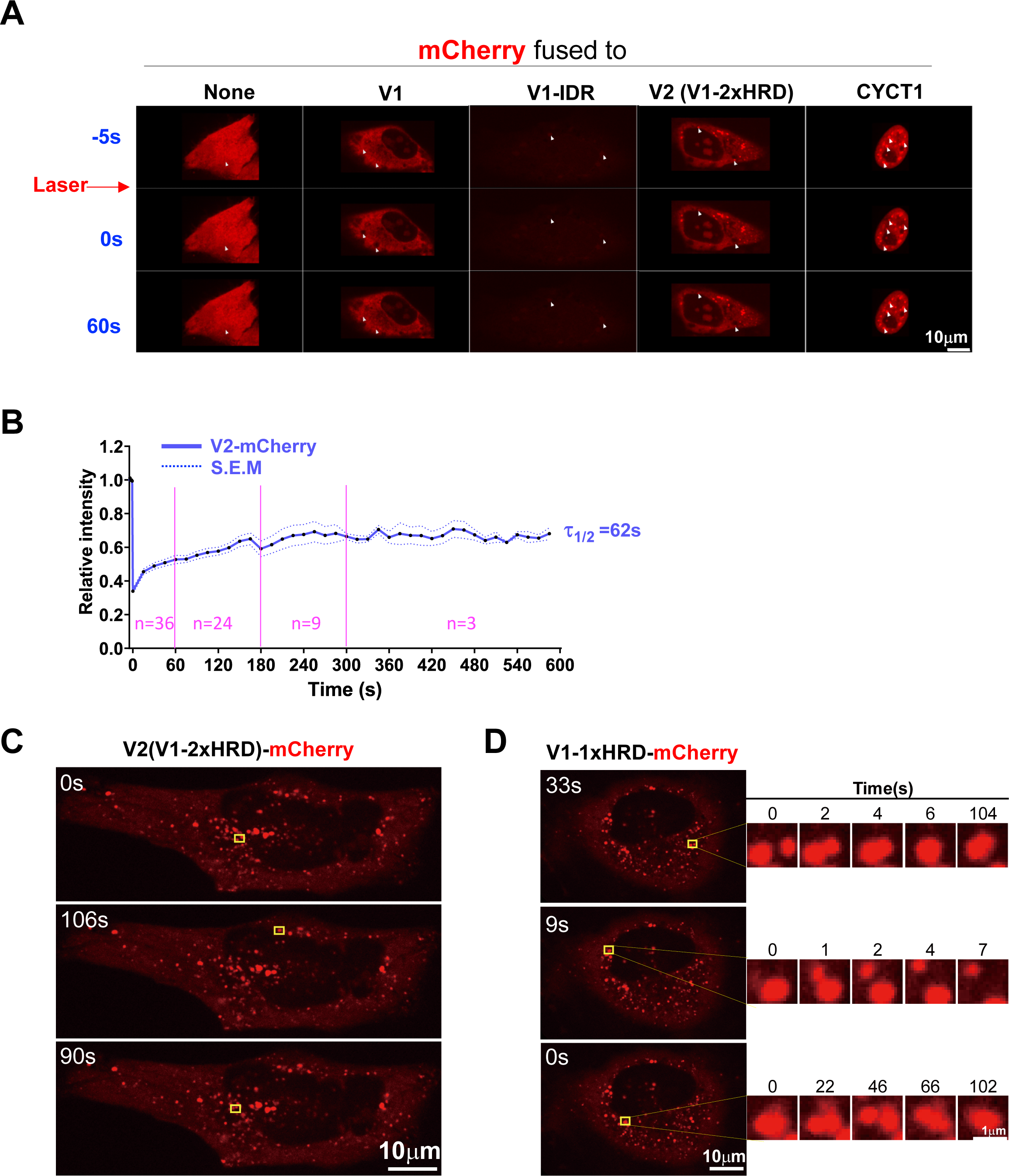
Characterization of the puncta formed by CYCT1 IDR and HRD. (A) Same as Fig. 4A, except that V1-IDR is not digitally enhanced. (B) FRAP analysis of V2-mCherry. Same as Fig. 4B except that fluorescence recovery was tracked for up to 10 min. Of note, the puncta tended to undergo fusion/fusion during this time; once this happened, the subsequent fluorescence recovery was no longer monitored or data discarded, which explains why the numbers of the puncta tracked decreased over time (from 36 to 3, pink). (C) Same as Fig. 4C, but showing the puncta (squares) in the context of the whole cell. The cell here was imaged at indicated times after video recording was started; these time points were set as time 0 in Fig. 4C. (D) The puncta formed by V1-1x HRD-mCherry also underwent fission/fusion.

**Fig. S6.**
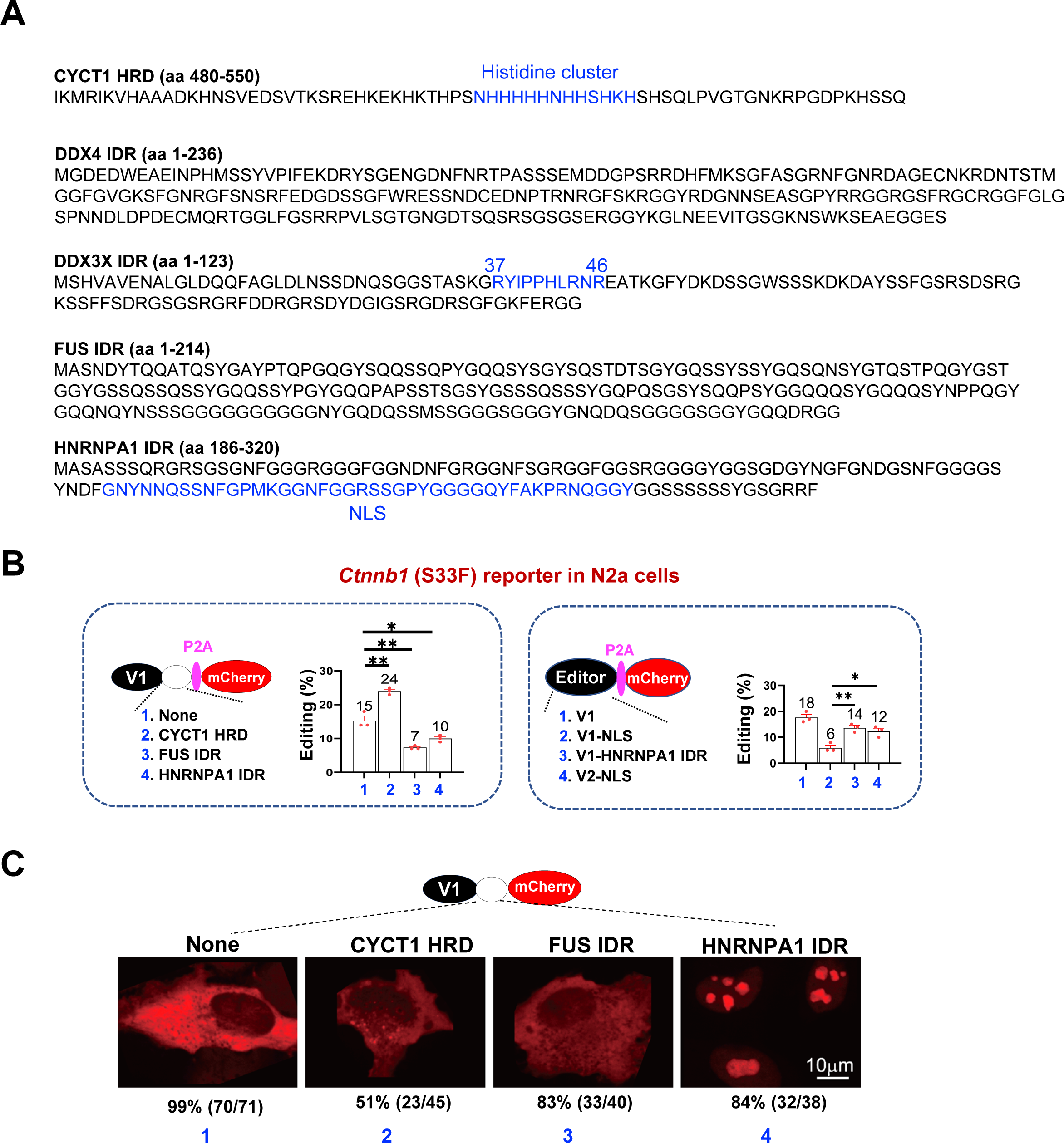
Effects of IDRs on V1. (A) Sequences of the IDRs tested in Fig. 5 and Fig. S5 B-C. Prominent features are highlighted in blue, including the 10-residue peptide in DDX3X IDR whose deletion compromises LLPS. (B) HNRNPA1 (but not FUS) IDR enhanced V1 editing. The IDRs were fused to V1 (left) or its nuclear version (right); nuclear localization was achieved by adding NLS (nuclear localization signal) to both termini of the editors. Same as Fig. 5B except for the IDRs used. (C) HNRNPA1 (but not FUS) IDR induced puncta formation by V1. Same as Fig. 5C except for the IDRs used.

**Fig. S7.**
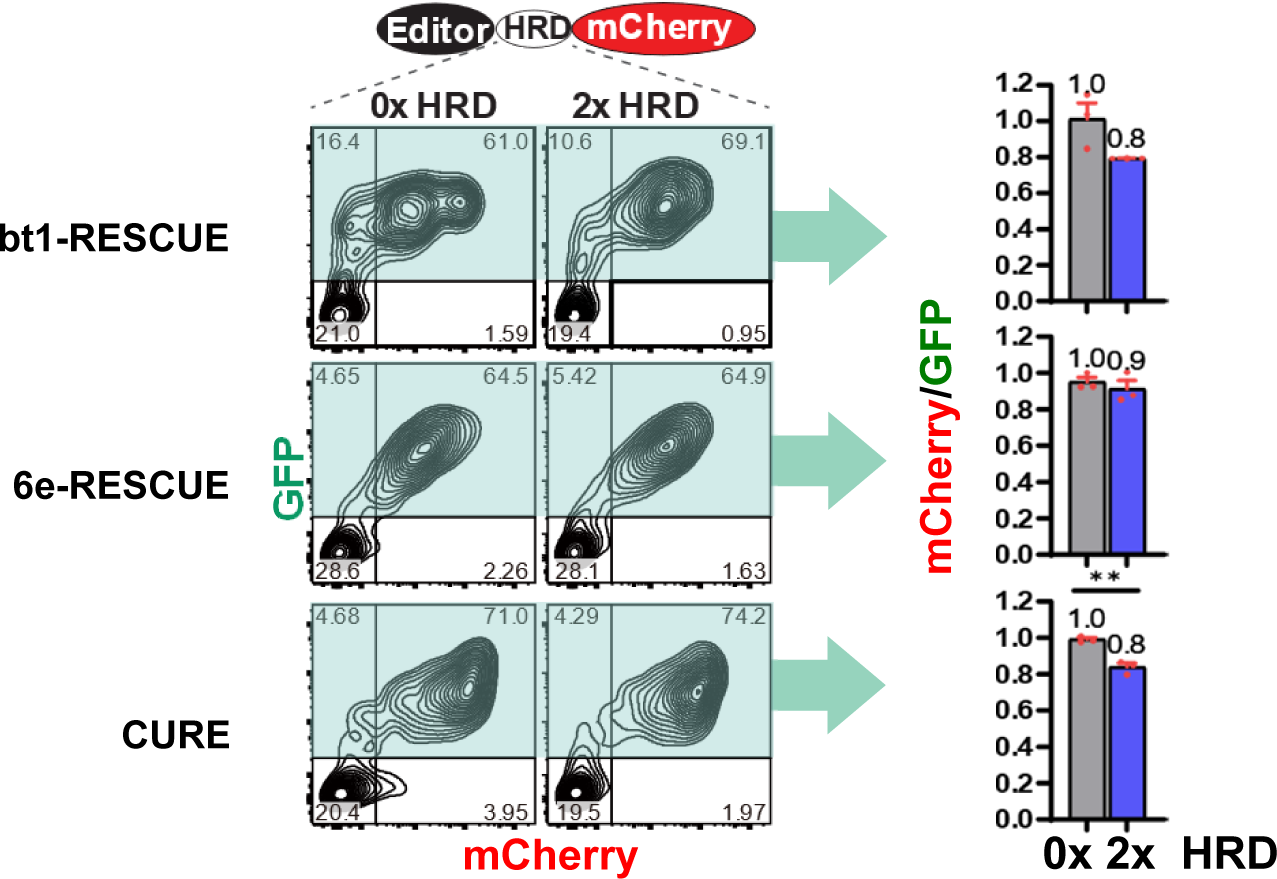
2x HRD does not markedly affect expression of indicated editors. The experiment was identical to that in Fig. 3A except that different editors were examined.

**Fig. S8.**
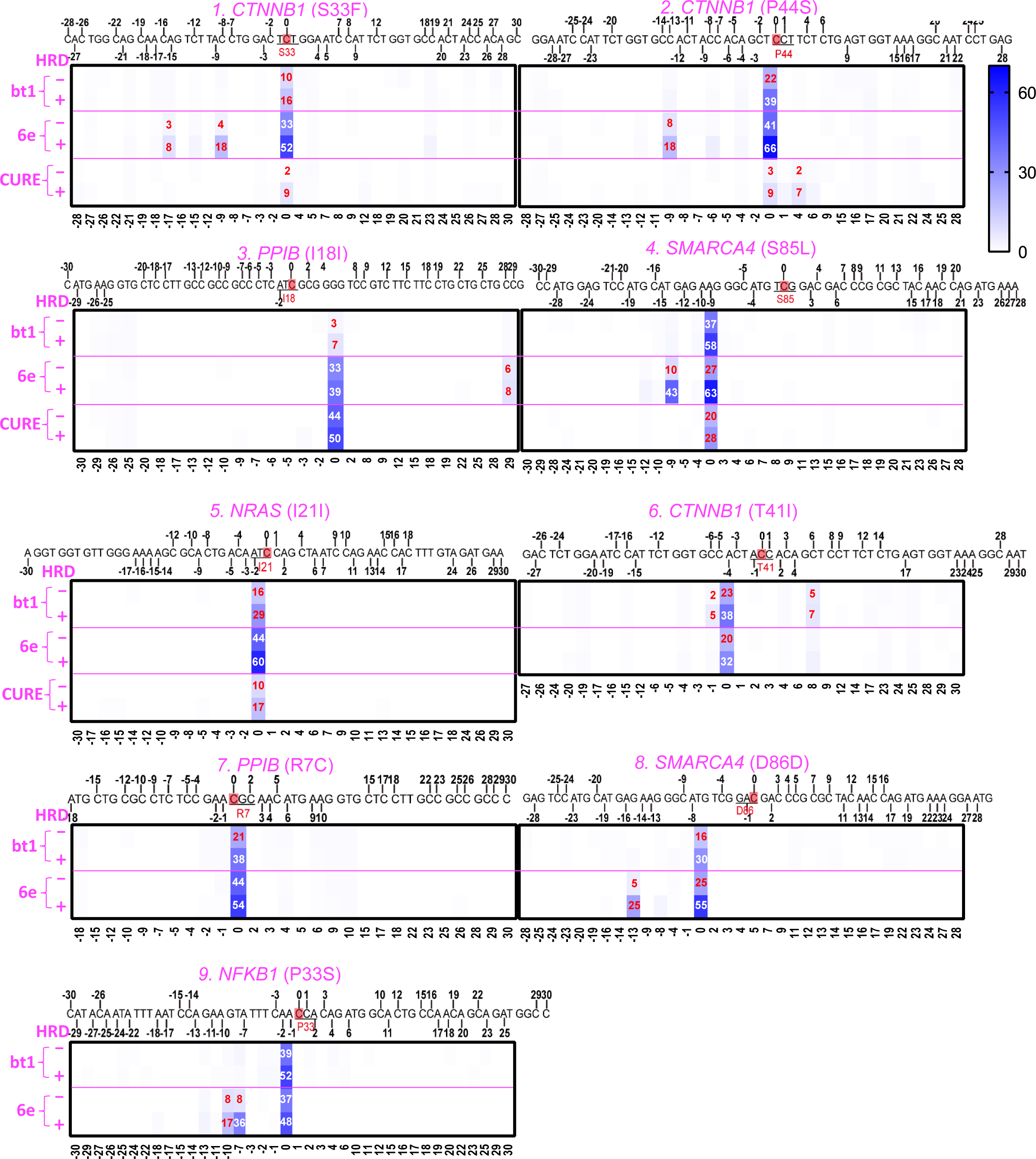
Effects of 2x HRD on bystander editing by three different editors. Same experiment as in Fig. 6A, but showing the targeted Cs (red) together with the bystanders (numbered bases) within the 60-nt flanking regions. The values in the heatmap denote editing rates, and are displayed only for the bases clearly and reproducibly edited. Note that 2x HRD did not exacerbate the bystander effect at Site 3, as in Fig. S4.

