## supplementary information for "Expanding RNA editing toolkit using an IDR-based strategy"

**1. Constructs**

**2×HRD protein sequence**

MGIINTLQKYYCRVRGGRCAVLSCLPKEEQIGKCSTRGRKCCRRKKGGSGGSGGSGGSGGSGGSMAVPETRPNHTIYINNLNSKIKKDELKKSLYAIFSQFGQILDILVPRQRTPRGQAFVIFKEVSSATNALRSMQGFPFYDKPMRIQYAKTDKRIPAKMKGTFVMHGSLQLPPLERLTLEFGGSGGSGGSGGSGGSGGSGSSQLHLPQVLADAVSRLVIGKFGDLTDNFSSPHARRIGLAGVVMTTGTDVKDAKVICVSTGAKCINGEYLSDRGLALNDCHAEIVSRRSLLRFLYTQLELYLNNEDDQKRSIFQKSERGGFRLKENIQFHLYISTSPCGDARIFSPHEAILEEPADRHPNRKARGQLRTKIEAGQGTIPVRNNASIQTWDGVLQGERLLTMSCSDKIARWNVVGIQGSLLSIFVEPIYFSSIILGSLYHGDHLSRAMYQRISNIEDLPPLYTLNKPLLTGISNAEARQPGKAPIFSVNWTVGDSAIEVINATTGKGELGRASRLCKHALYCRWMRVHGKVPSHLLRSKITKPNVYHETKLAAKEYQAAKARLFTAFIKAGLGAWVEKPTEQDQFSLTGSGSGSKMRIKVHAAADKHNSVEDSVTKSREHKEKHKTHPSNHHHHHNHHSHKHSHSQLPVGTGNKRPGDPKHSSQGSAAAGGSIKMRIKVHAAADKHNSVEDSVTKSREHKEKHKTHPSNHHHHHNHHSHKHSHSQLPVGTGNKRPGDPKHSSQGGSATNFSLLKQAGDVEENPGPMVSKGEEDNMAIIKEFMRFKVHMEGSVNGHEFEIEGEGEGRPYEGTQTAKLKVTKGGPLPFAWDILSPQFMYGSKAYVKHPADIPDYLKLSFPEGFKWERVMNFEDGGVVTVTQDSSLQDGEFIYKVKLRGTNFPSDGPVMQKKTMGWEASSERMYPEDGALKGEIKQRLKLKDGGHYDAEVKTTYKAKKPVQLPGAYNVNIKLDITSHNEDYTIVEQYERAEGRHSTGGMDELYK*

B-defensin TBP6.7 NES RESCUE HRD P2A mCherry

**IDR sequence**

MRIPVAGGDKAASSKPEEIKMRIKVHAAADKHNSVEDSVTKSREHKEKHKTHPSNHHHHHNHHSHKHSHSQLPVGTGNKRPGDPKHSSQTSNLAHKTYSLSSSFSSSSSTRKRGPSEETGGAVFDHPAKIAKSTKSSSLNFSFPSLPTMGQMPGHSSDTSGLSFSQPSCKTRVPHSKLDKGPTGANGHNTTQT

**gRNA expression vector**

GAGGGCCTATTTCCCATGATTCCTTCATATTTGCATATACGATACAAGGCTGTTAGAGAGATAATTGGAATTAATTTGACTGTAAACACAAAGATATTAGTACAAAATACGTGACGTAGAAAGTAATAATTTCTTGGGTAGTTTGCAGTTTTAAAATTATGTTTTAAAATGGACTATCATATGCTTACCGTAACTTGAAAGTATTTCGATTTCTTGGCTTTATATATCTTGTGGAAAGGACGAAACACCGGGCCAGATCTGAGCCTGGGAGCTCTCTGGCCTTATTGTCTTCGATATCGAAGACTTTTATT(GGCCAGATCTGAGCCTGGGAGCTCTCTGGCC)

U6 TAR hairpin	BbsI stuffer	(second optional TAR hairpin)

***Ctnnb1* reporter (CMV EGFP-SV40 intron-P2A-m*Ctnnb1*)**

ATGGTGAGCAAGGGCGAGGAGCTGTTCACCGGGGTGGTGCCCATCCTGGTCGAGCTGGACGGCGACGTAAACGGCCACAAGTTCAGCGTGTCCGGCGAGGGCGAGGGCGATGCCACCTACGGCAAGCTGACCCTGAAGTTCATCTGCACCACCGGCAAGCTGCCCGTGCCCTGGCCCACCCTCGTGACCACCCTGACCTACGGCGTGCAGTGCTTCAGCCGCTACCCCGACCACATGAAGCAGCACGACTTCTTCAAGTCCGCCATGCCCGAAGGCTACGTCCAGGAGCGCACCATCTTCTTCAAGGACGACGGCAACTACAAGACCCGCGCCGAGGTGAAGTTCGAGGGCGACACCCTGGTGAACCGCATCGAGCTGAAGGGCATCGACTTCAAGGAGGACGGCAACATCCTGGGGCACAAGCTGGAGTACAACTACAACAGCCACAACGTCTATATCATGGCCGACAAGCAGAAGAACGGCATCAAGGTGAACTTCAAGATCCGCCACAACATCGAGGACGGCAGCGTGCAGCTCGCCGACCACTACCAGCAGAACACCCCCATCGGCGACGGCCCCGTGCTGCTGCCCGACAACCACTACCTGAGCACCCAGTCCGCCCTGAGCAAAGACCCCAACGAGAAGCGCGATCACATGGTCCTGCTGGAGTTCGTGACCGCCGCCGGGATCACTCTCGGCATGGACGAGCTGTACAAGgtaagtttagtctttttgtcttttatttcaggtcccggatccggtggaaaggacgaaacaccgcaagtaaacccctaccaactggtcggggtttgaaaccatgcagacgcggttccactcgcagccacacaagtaaacccctaccaactggtcggggtttgaaactctagagtcggggcggcggtggtgcaaatcaaagaactgctcctcagtggatgttgcctttacttctagGGCTCCGCTACTAACTTCAGCCTGCTGAAGCAGGCTGGAGACGTGGAGGAGAACCCTGGACCTATGGCTACTCAAGCTGACCTGATGGAGTTGGACATGGCCATGGAGCCGGACAGAAAAGCTGCTGTCAGCCACTGGCAGCAGCAGTCTTACTTGGATTCTGGAATCCATTCTGGTGCCACCACCACAGCTCCTTCCCTGAGTGGCAAGGGCAACCCTGAGGAAGAAGATGTTGACACCTCCCAAGTCCTTTATGAATGGGAGCAAGGCTTTTCCCAGTCCTTCACGCAAGAGCAAGTAGCTGATATTGACGGGCAGTATGCAATGACTAGGGCTCAGAGGGTCCGAGCTGCCATGTTCCCTGAGACGCTAGATGAGGGCATGCAGATCCCATCCACGCAGTTTGACGCTGCTCATCCCACTAATGTCCAGCGCTTGGCTGAACCATCACAGATGTTGAAACATGCAGTTGTCAATTTGATTAACTATCAGGATGACGCGGAACTTGCCACACGTGCAATTCCTGAGCTGACAAAACTGCTAAACGATGAGGACCAGGTGGTAGTTAATAAAGCTGCTGTTATGGTCCATCAGCTTTCCAAAAAGGAAGCTTCCAGACATGCCATCATGCGCTCCCCTCAGATGGTGTCTGCCATTGTACGCACCATGCAGAATACAAATGATGTAGAGACAGCTCGTTGTACTGCTGGGACTCTGCACAACCTTTCTCACCACCGCGAGGGCTTGCTGGCCATCTTTAAGTCTGGTGGCATCCCAGCGCTGGTGAAAATGCTTGGGTCACCAGTGGATTCTGTACTGTTCTACGCCATCACGACACTGCATAATCTCCTGCTCCATCAGGAAGGAGCTAAAATGGCAGTGCGCCTAGCTGGTGGACTGCAGAAAATGGTTGCTTTGCTCAACAAAACAAACGTGAAATTCTTGGCTATTACAACAGACTGCCTTCAGATCTTAGCTTATGGCAATCAAGAGAGCAAGCTCATCATTCTGGCCAGTGGTGGACCCCAAGCCTTAGTAAACATAATGAGGACCTACACTTATGAGAAGCTTCTGTGGACCACAAGCAGAGTGCTGAAGGTGCTGTCTGTCTGCTCTAGCAACAAGCCGGCCATTGTAGAAGCTGGTGGGATGCAGGCACTGGGGCTTCATCTGACAGACCCAAGTCAGCGACTTGTTCAAAACTGTCTTTGGACTCTCAGAAACCTTTCAGATGCAGCGACTAAGCAGGAAGGGATGGAAGGCCTCCTTGGGACTCTAGTGCAGCTTCTGGGTTCCGATGATATAAATGTGGTCACCTGTGCAGCTGGAATTCTCTCTAACCTCACTTGCAATAATTACAAAAACAAGATGATGGTGTGCCAAGTGGGTGGCATAGAGGCTCTTGTACGCACCGTCCTTCGTGCTGGTGACAGGGAAGACATCACTGAGCCTGCCATCTGTGCTCTTCGTCATCTGACCAGCCGGCATCAGGAAGCCGAGATGGCCCAGAATGCCGTTCGCCTTCATTATGGACTGCCTGTTGTGGTTAAACTCCTGCACCCACCATCCCACTGGCCTCTGATAAAGGCAACTGTTGGATTGATTCGAAACCTTGCCCTTTGCCCAGCAAATCATGCGCCTTTGCGGGAACAGGGTGCTATTCCACGACTAGTTCAGCTGCTTGTACGAGCACATCAGGACACCCAACGGCGCACCTCCATGGGTGGAACGCAGCAGCAGTTTGTGGAGGGCGTGCGCATGGAGGAGATAGTAGAAGGGTGTACTGGAGCTCTCCACATCCTTGCTCGGGACGTTCACAACCGGATTGTAATCCGAGGACTCAATACCATTCCATTGTTTGTGCAGTTGCTTTATTCTCCCATTGAAAATATCCAAAGAGTAGCTGCAGGGGTCCTCTGTGAACTTGCTCAGGACAAGGAGGCTGCAGAGGCCATTGAAGCTGAGGGAGCCACAGCTCCCCTGACAGAGTTACTCCACTCCAGGAATGAAGGCGTGGCAACATACGCAGCTGCTGTCCTATTCCGAATGTCTGAGGACAAGCCACAGGATTACAAGAAGCGGCTTTCAGTCGAGCTGACCAGTTCCCTCTTCAGGACAGAGCCAATGGCTTGGAATGAGACTGCAGATCTTGGACTGGACATTGGTGCCCAGGGAGAAGCCCTTGGATATCGCCAGGATGATCCCAGCTACCGTTCTTTTCACTCTGGTGGATACGGCCAGGATGCCTTGGGGATGGACCCTATGATGGAGCATGAGATGGGTGGCCACCACCCTGGTGCTGACTATCCAGTTGATGGGCTGCCTGATCTGGGACACGCCCAGGACCTCATGGATGGGCTGCCCCCAGGTGATAGCAATCAGCTGGCCTGGTTTGATACTGACCTGTAAATCGTCCTTTAGGTAAGAAAGCTTATAAAAGCCAGTGTGGGTGAATACTTTACTCTGCCTGCAGAACTCCAGAAAGACTTGGTAGGGTGGGAATGGTTTTAGGCCTGTTTGTAAATCTGCCACCAAACAGATACATACCTTGGAAGGAGATGTTCATGTGTGGAAGTTTCTCACGTTGATGTTTTTGCCACAGCTTTTGCAGCGTTATACTCAGATGAGTAACATTTGCTGTTTTCAACATTAATAGCAGCCTTTCTCTCTATACAGCTGTAGTGTCTGAACGTGCATTGTGATTGGCCTGTAGAGTTGCTGAGAGGGCTCGAGGGGTGGGCTGGTATCTCAGAAAGTGCCTGACACACTAACCAAGCTGAGTTTCCTATGGGAACAGTCGAAGTACGCTTTTTGTTCTGGTCCTTTTTGGTCGAGGAGTAACAATACAAATGGATTTGGGGAGTGACTCACGCAGTGAAGAATGCACACGAATGGATCACAAGATGGCGTTATCAAACCCTAGCCTTGCTTGTTCTTTGTTTTAATATCTGTAGTGGTGCTGACTTTGCTTGCTTTTATTTTTTGCAGTAACTGTTAGTTTTTAAGTAGTGTTATGTTCTAGTGAACCTGCTACAGCAATTTCTGATTTCTAAGAACCGAGTAATGGTGTAGAACACTAATTCATAATCACGCTAATTGTAATCTGGAGACGTGTAACATTGTGTAGCCTTTTGTATAAATAGACAGATAGAAATGGTCCGATTAGTTTCCTTTTTAATATGCTTAAAATAAGCAGGTGGATCTATTTCATGTTTTTGAACAAAAACTTTATCGGGGATACGTGCGGTAGGGTAAATCAGTAAGAGGTGTTATTTGAGCCTTGTTTTGGACAGTATACCAGTTGCCTTTTATCCCAAAGTTGTTGTAACCTGCTGTGATACAATGCTTCAACAGATGCGGTTATAGAAATGGTTCAGAATTAAACTTTTAATTCATTCAAAA

EGFP	SV40 intron P2A *Ctnnb1* cDNA

**2. gRNAs (mismatched base capitalized)**

**gRNA for C-RESCUE**

hCTNNB1 S33F C-flip

30/22 ggattccaCagtccaggtaagactgttgct

hCTNNB1 P44S C-flip

30/22 cagagaagCagctgtggtagtggcaccaga

hCTNNB1 T41I C-flip

30/22 gagctgtgCtagtggcaccagaatggattc

hPPIB I18I C-flip

30/22 gaccccgcCatgagggcggcggcaaggagc

hPPIB R7C C-flip

30/22 catgttgcCttcggagaggcgcagcatcca

hSMACA4 S85L C-flip

30/22 ggtcgtccCacatgcccttctcatgcatgg

hSMACA4 D86D C-flip

30/22 cgcgggtcCtccgacatgcccttctcatgc

hNRAS I21I C-flip

30/22 attagctgCattgtcagtgcgcttttccca

hNKFB1 P33S C-flip

30/22 catctgtgCttgaaatacttctggattaaa

mCTNNB1 S33F C-flip

20/10 atggattccaCaatccaagt

20/15 ttccaCaatccaagtaagac

30/5	ggtggtggcaccagaatggattccaCaatc

30/10 tggcaccagaatggattccaCaatccaagt

30/15 ccagaatggattccaCaatccaagtaagac

30/20 atggattccaCaatccaagtaagactgctg

30/22 ggattccaCaatccaagtaagactgctgct

30/24 attccacaatccaagtaagactgctgctgc

30/25 ttccaCaatccaagtaagactgctgctgcc

30/26 tccaCaatccaagtaagactgctgctgcca

40/20 tggcaccagaatggattccaCaatccaagtaagactgctg

50/25 ggtggtggcaccagaatggattccaCaatccaagtaagactgctgctgcc

50/30 tggcaccagaatggattccaCaatccaagtaagactgctgctgccagtgg

mCTNNB1 S33F U-flip

30/5	ggtggtggcaccagaatggattccaUaatc

30/10 tggcaccagaatggattccaUaatccaagt

30/15 ccagaatggattccaUaatccaagtaagac

30/20 atggattccaUaatccaagtaagactgctg

30/22 ggattccaUaatccaagtaagactgctgct

30/24 attccaUaatccaagtaagactgctgctgc

30/25 ttccaUaatccaagtaagactgctgctgcc

30/26 tccaUaatccaagtaagactgctgctgcca

mCTNNB1 S37F U-flip

30/15 gtggtggtggcaccaUaatggattccagaa

30/20 ggtggcaccaUaatggattccagaatccaa

30/22 tggcaccaUaatggattccagaatccaagt

mCTNNB1 T41I U-flip

30/15 agggaaggagctgtgUtggtggcaccagaa

30/20 aggagctgtgUtggtggcaccagaatggat

30/22 gagctgtgUtggtggcaccagaatggattc

mCTNNB1 S45F U-flip

30/15 cccttgccactcaggUaaggagctgtggtg

30/20 gccactcaggUaaggagctgtggtggtggc

30/22 cactcaggUaaggagctgtggtggtggcac

**gRNA for dCas13bt1-RESCUE**

hCTNNB1 S33F U-flip

30/22 ggattccaUagtccaggtaagactgttgct

hCTNNB1 T41I U-flip

30/22 gagctgtgUtagtggcaccagaatggattc

hCTNNB1 P44S U-flip

30/22 cagagaagUagctgtggtagtggcaccaga

hPPIB I18I U-flip

30/22 gaccccgcUatgagggcggcggcaaggagc

hPPIB R7C U-flip

30/22 catgttgcUttcggagaggcgcagcatcca

hNKFB1 P33S U-flip

30/22 catctgtgUttgaaatacttctggattaaa

hSMARCA4 S85L U-flip

30/22 ggtcgtccUacatgcccttctcatgcatgg

hSMARCA4 D86D U-flip

30/22 cgcgggtcUtccgacatgcccttctcatgc

hNRAS I21I U-flip

30/22 attagctgUattgtcagtgcgcttttccca

**gRNA for dCas6e-RESCUE**

hCTNNB1 S33F C-flip

30/26 tccaCagtccaggtaagactgttgctgcca

hCTNNB1 T41I C-flip

30/26 tgtgCtagtggcaccagaatggattccaga

hCTNNB1 P44S C-flip

30/26 gaagCagctgtggtagtggcaccagaatgg

hPPIB I18I C-flip

30/26 ccgcCatgagggcggcggcaaggagcacct

hPPIB R7C C-flip

30/26 ttgcCttcggagaggcgcagcatccacagg

hNKFB1 P33S C-flip

30/26 tgtgCttgaaatacttctggattaaatatt

hSMARCA4 S85L C-flip

30/26 gtccCacatgcccttctcatgcatggactc

hSMARCA4 D86D C-flip

30/26 ggtcCtccgacatgcccttctcatgcatgg

hNRAS I21I C-flip

30/26 gctgCattgtcagtgcgcttttcccaacac

**gRNA for CURE**

hCTNNB1 S33F

S32L14 gtggcaccagaatggataagactgttgctgcc

hCTNNB1 T41I

S32L14 ccactcagagaaggagccagaatggattccag

hCTNNB1 P44S

S32L14 tgcctttaccactcagtagtggcaccagaatg

hPPIB I18I

S32L14 caggaagaagacggacggcggcaaggagcacc

hPPIB R7C

S32L14 caaggagcaccttcatggcgcagcatccacag

hNKFB1 P33S

S32L14 ctgttggcagtgccatcttctggattaaatat

hSMARCA4 S85L

S32L14 tggttgtagcgcgggtttctcatgcatggact

hSMARCA4 D86D

S32L14 catctggttgtagcgcgcccttctcatgcatg

hNRAS I21I

S32L14 aaagtggttctggatttgcgcttttcccaaca

**PCR primers for amplifying cDNA before NGS**

mCTNNB1 reporter forward: TCTCGGCATGGACGAGCTGTACAAGGGCTCC

mCTNNB1 reporter reverse: AACATGGCAGCTCGGACCCTC

mCTNNB1 forward: CTGCGTGGACAATGGCTACTCAAGCTGAC

mCTNNB1 reverse: AACATGGCAGCTCGGACCCTC

hCTNNB1 forward: GAGAACCCTGGACCTAAGCTGATTTGATGGAGTTGGAC

hCTNNB1 reverse: GCCCCTAACCGGTTTAAAGGACTGAGAAAATCCCTGT

hPPIB forward: GAGAACCCTGGACCTCTGCTTTCGCCTCCGCCTG

hPPIB reverse: GCCCCTAACCGGTTTAGCTGTTTTTGTAGCCAAATCCTTT

hSMACA4 forward: CCTCAGGACAACATGCACCA

hSMACA4 reverse: GCTGGAACTGGACTAGAGGC

hNRAS forward: TCTTGCTGGTGTGAAATGACTGA

hNRAS reverse: CTCTCATGGCACTGTACTCT

hNFKB1 forward: TGGGAAGGCCTGAACAAATG

hNFKB1 reverse: TGTTCTTTTCACTAGAGGCACCA
